# Co-Targeting Nuclear Export and Translation Initiation Uncovers a Therapeutic Vulnerability in Lethal Prostate Cancer

**DOI:** 10.64898/2026.05.04.722693

**Authors:** Jessica D. Kindrick, Kinjal Bhadresha, Xiaohu Zhang, Erica L. Beatson, Spencer S. Gaut, Benjamin C. Brim, Patrick J. Signorelli, Roger Depaz, Jessica L. Horner, Posey S. Whidden, John M. Ching, Kelli M. Wilson, Savannah Wood, Crystal McKnight, Erin Beck, Carleen Klumpp-Thomas, Ross Lake, Elijah Edmondson, Michele Ceribelli, Cindy H. Chau, Craig J. Thomas, William D Figg

**Affiliations:** Genitourinary Malignancies Branch, Center for Cancer Research, National Cancer Institute, National Institutes of Health, Bethesda, MD, USA; Division of Preclinical Innovation, National Center for Advancing Translational Sciences, National Institutes of Health, Rockville, MD, USA; Clinical Pharmacology Program, Office of the Clinical Director, Center for Cancer Research, National Cancer Institute, Bethesda, Maryland, USA; Laboratory of Cancer Biology and Genetics, Center for Cancer Research, National Cancer Institute, National Institutes of Health, Bethesda, MD, USA; Molecular Histopathology Lab, Laboratory of Animal Sciences Program, Frederick National Lab for Cancer Research

## Abstract

Metastatic castration-resistant prostate cancer (mCRPC) remains lethal as adaptive resistance to standard-of-care therapy develops, often driven by AR splice variants alongside transcriptional and translational reprogramming. To identify strategies capable of overcoming these mechanisms, we performed an unbiased high-throughput screen of 2,480 mechanistically annotated compounds across advanced prostate cancer models. Exportin-1 (XPO1)-mediated nuclear export emerged as a critical dependency, and matrix-based combination screening uncovered robust synergy between inhibitors of XPO1 and the translation initiation factor EIF4A1. Dual inhibition induced coordinated disruption of oncogenic protein networks, including AR/AR-V7, triggering apoptosis and suppressing cell-cycle and metabolic programs. These effects extended to genetically diverse patient-derived organoids and in vivo xenografts at low doses, approximately 8-fold (Eltanexor) and 12-fold (Zotatifin) below established human single-agent regimens. Together, these findings reveal concurrent control of nuclear export and protein translation as a therapeutic vulnerability in mCRPC, providing a strong rationale for clinical evaluation of XPO1-EIF4A1 co-inhibition to overcome AR-driven resistance.

**STATEMENT OF SIGNIFICANCE:** Unbiased combinatorial screening reveals co-inhibition of nuclear export and translation initiation as a vulnerability in metastatic castration-resistant prostate cancer. Dual targeting of XPO1 and EIF4A1 drives synergistic collapse of oncogenic protein networks, including AR/AR-V7 signaling, to overcome key resistance mechanisms and induce potent antitumor responses across heterogeneous models. Notably, these effects are achieved at substantially reduced doses using clinically tractable agents, defining a mechanistically grounded therapeutic strategy poised for rapid clinical translation.

## INTRODUCTION

Metastatic castration-resistant prostate cancer (mCRPC) remains a lethal disease state for which durable therapeutic options are limited.^1,2^ Despite substantial advances in androgen receptor (AR)-directed therapies, resistance is nearly universal and is frequently driven by AR splice variants, lineage plasticity, and extensive rewiring of transcriptional and translational programs.^3–5^ Together, these adaptations result in transient clinical responses followed by inevitable disease progression in heavily pretreated patients. These features highlight a fundamental challenge in advanced prostate cancer: targeting a single dominant pathway is often insufficient to achieve durable disease control across heterogeneous tumors.^6^ Consequently, there is a pressing need for therapeutic strategies that move beyond incremental modifications of existing regimens and instead leverage coordinated disruption of multiple oncogenic processes.

One effective strategy to overcome therapeutic resistance is the use of rational drug combinations that simultaneously target complementary vulnerabilities. In principle, combinations composed of agents with distinct mechanisms of action may reduce the likelihood of adaptive resistance while limiting overlapping toxicities, particularly when synergistic interactions permit dose reduction. However, in mCRPC, most combination regimens have been assembled empirically around existing standards of care rather than discovered through systematic, unbiased approaches, and many have failed to produce meaningful or durable clinical benefit.^7^ These outcomes underscore a broader limitation in current combination strategies: without a framework to identify synergistic and biologically coherent interactions, many potentially effective combinations, particularly those targeting noncanonical or orthogonal processes, remain unexplored.

Phenotypic screening of mechanistically annotated drug libraries provides a powerful alternative framework to uncover therapeutically actionable interactions without presupposing a specific target or pathway.^8^ Unlike hypothesis-driven combination testing, this approach enables systematic evaluation of drug activity and synergy across diverse concentration ranges, temporal windows, and disease contexts, while capturing emergent phenotypes that reflect integrated cellular responses. Importantly, when implemented quantitatively, phenotypic screening can prioritize combinations that achieve efficacy at clinically relevant exposures and reveal mechanistic dependencies that are not apparent from single agent analyses alone.

In the present study, we applied a robotically automated, quantitative high-throughput screening platform to interrogate 2,480 approved and investigational compounds across multiple prostate cancer models. This platform enables high-resolution single agent profiling followed by structured all-versus-all matrix combination testing, allowing unbiased identification of synergistic interactions that suppress viability and induce apoptosis at clinically achievable doses.^9–11^ Using this framework, we sought to identify combination strategies capable of overcoming key resistance features of advanced prostate cancer, including AR splice variant AR-V7-driven signaling, while maintaining translational feasibility.

Here, we identify coordinated inhibition of nuclear export and translation initiation as a highly effective combination strategy in prostate cancer. Through unbiased screening, we uncover strong and reproducible synergy between inhibitors of exportin-1 (XPO1) and the translation initiation factor EIF4A1, which are two fundamental processes that converge on oncogenic protein homeostasis but have not previously been co-targeted in this disease context. We demonstrate that this combination suppresses core oncogenic protein networks, induces apoptosis at reduced doses, and produces robust antitumor activity across genetically and phenotypically diverse in vitro and in vivo models, supporting its advancement toward clinical evaluation in patients with advanced prostate cancer.

## RESULTS

### Quantitative high-throughput screening identifies nuclear export (XPO1) as a therapeutic vulnerability in prostate cancer

To identify therapeutic vulnerabilities in prostate cancer, we performed a quantitative high-throughput screen (qHTS) using an oncology-focused small-molecule library (n = 2,480 compounds) across AR-V7-expressing LNCaP-95 and VCaP-CR cells (Figure 1A). Responses were assessed using CellTiter-Glo (CTG) luminescent cell viability readouts at 48 h in LNCaP-95 cells and 72 h in VCaP-CR cells. Compounds produced a broad range of effects on cell viability, with high concordance between the two models (Pearson R = 0.72, p < 2.2 × 10^-16^), demonstrating the robustness of the screening platform (Figure 1B). Integrating responses across cell lines highlighted both highly active and inactive agents, with XPO1 inhibitors among the most potent (Figure 1C, Table S1). Drug Target Set Enrichment Analysis confirmed statistically significant enrichment of XPO1-targeting compounds among the top-ranking hits, implicating nuclear export as a dependency in AR-V7-expressing prostate cancer (Figure 1D).

**Figure 1.**
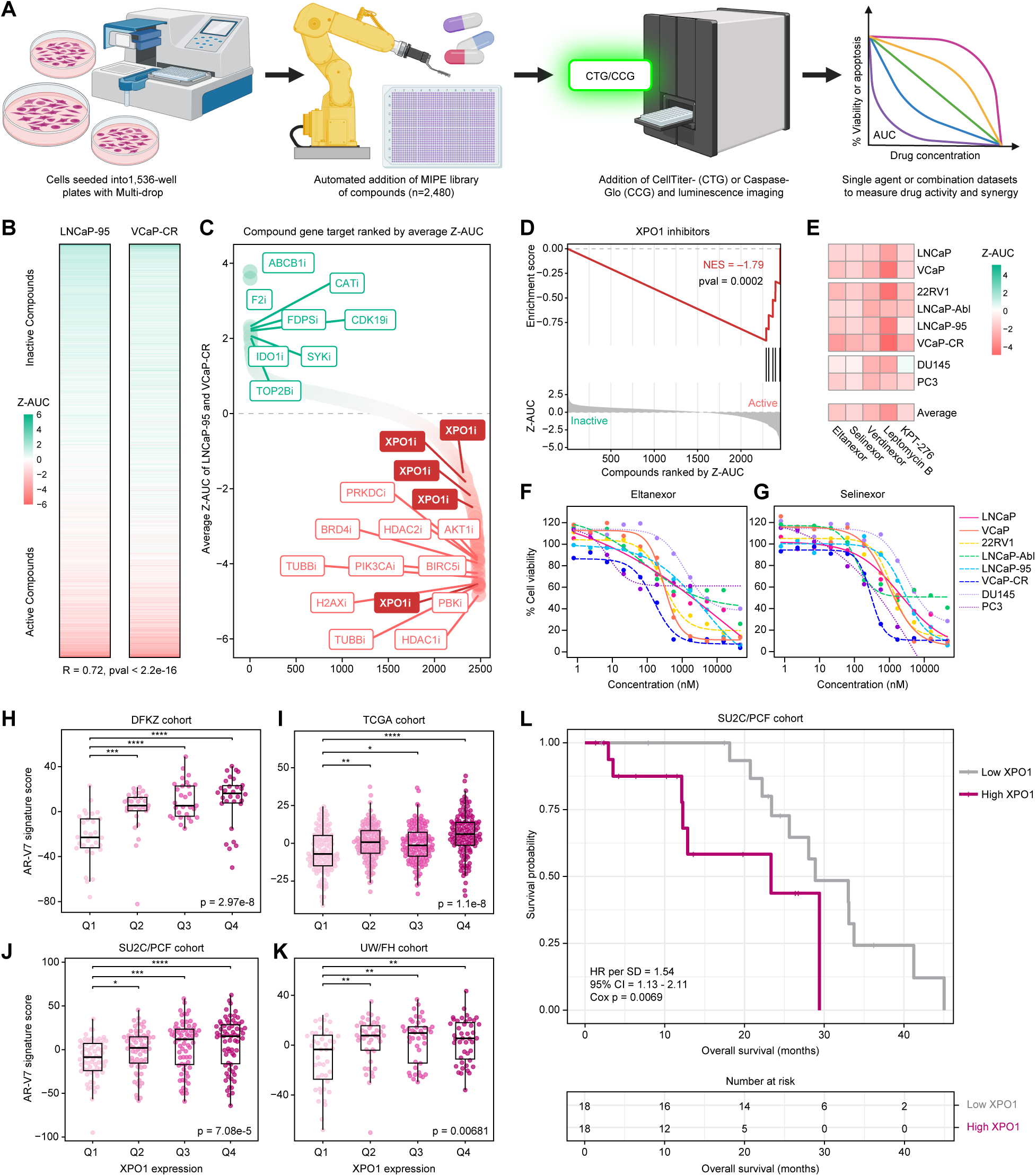
Quantitative high-throughput screening identifies nuclear export (XPO1) as a therapeutic vulnerability in prostate cancer. (A) Schematic of the quantitative high-throughput screening (qHTS) workflow using the Mechanism Interrogation PlatE (MIPE) small-molecule library to measure drug activity and synergy. Created with Biorender.com. (B) Heatmap of Z-normalized area under the curve (Z-AUC) values derived from 11-point dose-response viability measurements (CellTiter-Glo) in AR-V7-expressing LNCaP-95 (48 h readout) and VCaP-CR (72 h readout) cells, with the Pearson correlation coefficient between cell types indicated. (C) Mean Z-AUC values across LNCaP-95 and VCaP-CR cells for each compound, ranked in descending order and annotated by primary molecular target (“i” denotes inhibitor). (D) Drug Target Set Enrichment Analysis (DSEA) demonstrating significant enrichment of XPO1-targeting compounds among the most active agents when compounds are ranked by Z-AUC. (E) Heatmap of Z-AUC values for XPO1 inhibitors (Eltanexor, Selinexor, Leptomycin B, and KPT-276) across the indicated panel of prostate cancer cells (LNCaP, VCaP, 22Rv1, LNCaP-Abl, LNCaP-95, VCaP-CR, DU145, and PC3). (F) Single agent dose-response curves displaying percent cell viability following Eltanexor treatment across an 11-point concentration range (0.8 nM-45 μM; 1:3 dilution series) in representative castration-sensitive, castration-resistant, and aggressive variant prostate cancer cells. (G) As in (F), dose-response curves for Selinexor. (H-K) Clinical prostate cancer samples were stratified into quartiles based on XPO1 expression, ranging from low (Q1) to high (Q4). An AR-V7 target gene score was calculated as the summed expression of genes from the Sharp et al. (2019) signature (n = 59). Differences across quartiles were assessed using the Kruskal-Wallis test, followed by Dunn’s post hoc testing with Benjamini-Hochberg correction. Analyses were performed in primary prostate cancer cohorts from (H) DKFZ and (I) TCGA, and metastatic castration-resistant prostate cancer cohorts from (J) SU2C/PCF and (K) UW/FH. Statistical significance is indicated (*p < 0.05, **p < 0.01, ***p < 0.001, ****p < 0.0001). (L) Kaplan-Meier overall survival analysis in the SU2C/PCF cohort comparing low (Q1) versus high (Q4) XPO1 expression. Hazard ratio (HR) per standard deviation (SD) with 95% confidence interval (CI) and Cox proportional hazards p-value are shown; numbers at risk are indicated below. See also Figure S1.

We next expanded XPO1 inhibitor testing across a broader panel of 8 prostate cancer models (LNCaP, VCaP, 22Rv1, LNCaP-95, VCaP-CR, DU145, and PC3), including castration-sensitive (CSPC), castration-resistant (CRPC), and aggressive variant prostate cancer (AVPC) cell lines. All five XPO1 inhibitors tested (Eltanexor, Selinexor, Verdinexor, Leptomycin B, and KPT-276) demonstrated potent reductions in cell viability across these models, with Eltanexor and Selinexor representing the most clinically advanced and translationally relevant compounds (Figure 1E-G, Figure S1).

Given previous reports that XPO1 inhibition can sequester AR-V7 mRNA in the nucleus and reduce its transcriptional activity^12^, we hypothesized that XPO1 expression might correlate with AR-V7 target gene activity in clinical prostate cancer specimens. Consistent with this, tumors with higher XPO1 expression exhibited increased AR-V7 transcriptional activity across primary (DKFZ, TCGA) and metastatic (SU2C/PCF, UW/FH) cohorts (Figure 1H-K).^13^ Elevated XPO1 expression was also associated with decreased overall survival in the metastatic castration-resistant SU2C/PCF cohort (HR = 1.54 per SD, 95% CI 1.13-2.11, Cox p = 0.0069; Figure 1L), supporting XPO1 as a clinically relevant therapeutic target.

### Matrix combination screening reveals robust synergy between XPO1 and EIF4A1 inhibition

To identify synergistic drug combinations in AR-V7-expressing prostate cancer, we selected 42 compounds based on single agent activity (Figure 1) and mechanistic interest to test all possible pairwise interactions (n = 861). Single agent dose-response data were used to define appropriate concentration ranges for each compound, and each drug pair was evaluated using a 10 × 10 matrix. Responses were again assessed by CTG viability readouts at 48 h in LNCaP-95 cells and at 72 h in VCaP-CR cells. Excess Highest Single Agent (Excess HSA) scores were calculated for each matrix to quantify synergistic and antagonistic interactions, and values were averaged across both cell lines (Table S2).

Unsupervised hierarchical clustering of compounds based on correlation of Excess HSA profiles revealed mechanistically related clusters with highly similar combination behaviors (Figure 2A), indicating that compounds sharing mechanistic targets exhibit conserved interaction patterns across diverse partners. Notably, the XPO1 inhibitors Eltanexor and Selinexor clustered together, reflecting highly similar combination profiles. Additional clusters included AKT inhibitors (AZD-5363 and Ipatasertib), BCL2 inhibitors (Navitoclax and Venetoclax), and multiple tubulin-targeting agents (Docetaxel, Cabazitaxel, Vincristine sulfate, and PTC-596), demonstrating that dual-drug combination response patterns reflect underlying biological relationships.

**Figure 2.**
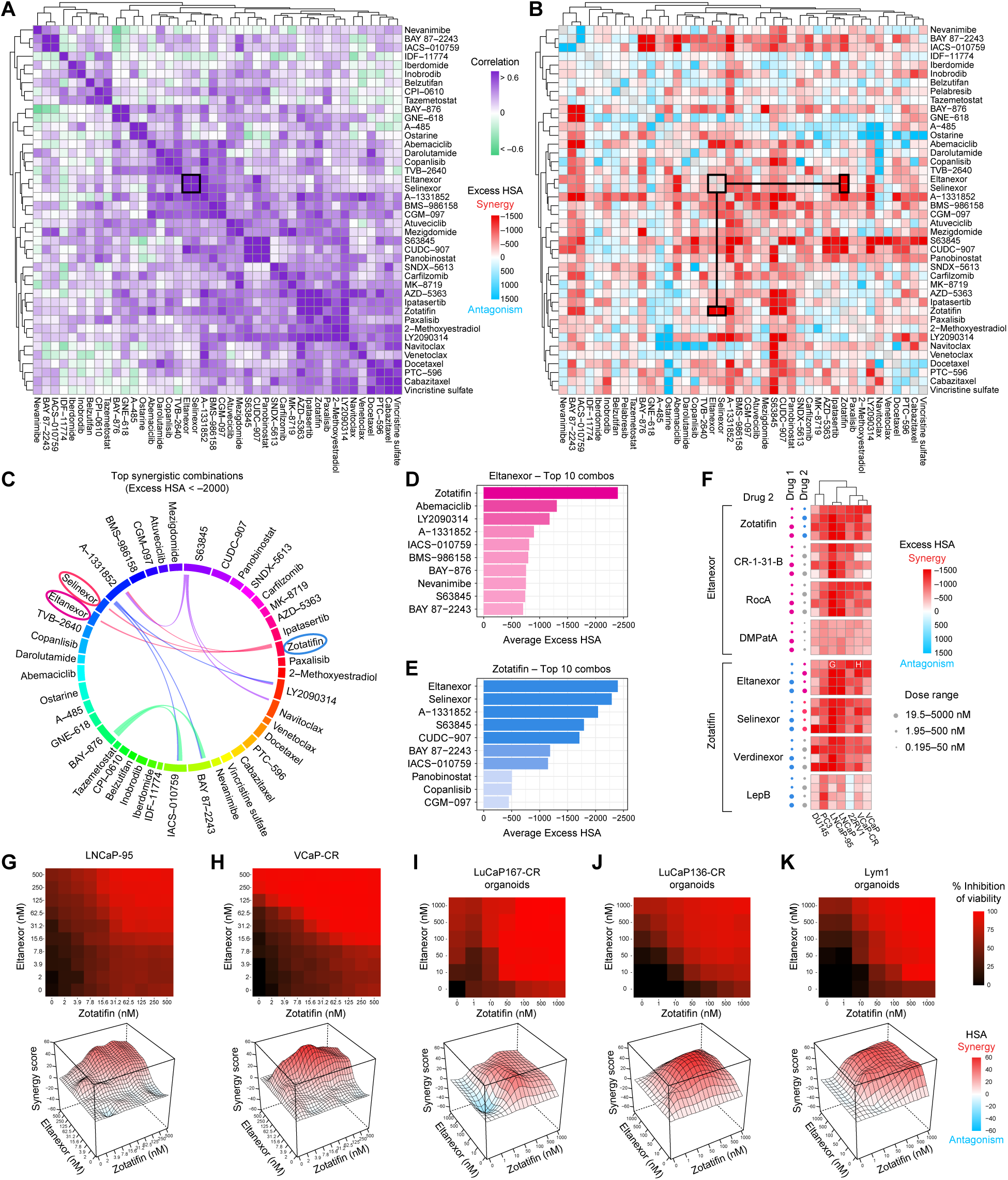
High-throughput combination drug screening reveals robust synergy between XPO1 and EIF4A1 inhibition. (A) Unsupervised hierarchical clustering of a 42-drug all-versus-all combination screen based on correlation of Excess Highest Single Agent (Excess HSA) scores, identifying clusters of compounds with similar combination profiles. Compounds selected based on single agent activity and mechanistic relevance were tested across 10 × 10 dose matrices (n = 861 combinations) in LNCaP-95 and VCaP-CR cells, with Excess HSA values averaged across both models. The black box highlights clustering of combination profiles involving the XPO1 inhibitors Eltanexor and Selinexor. (B) Excess HSA synergy scores overlaid onto the same unsupervised clustering heatmap, highlighting synergistic (red) and antagonistic (blue) interactions. The black boxes highlight strong synergy between XPO1 inhibitors (Eltanexor, Selinexor) and the EIF4A1 inhibitor Zotatifin. (C) Circos plot displaying the most synergistic drug combinations (Excess HSA < -2000) identified in the screen. (D) Top 10 most synergistic combinations with Eltanexor, ranked by mean Excess HSA across LNCaP-95 and VCaP-CR cells. (E) Top 10 most synergistic combinations with Zotatifin, ranked by mean Excess HSA across LNCaP-95 and VCaP-CR cells. (F) Targeted combination screening evaluating XPO1 and EIF4A1 inhibition across prostate cancer models. Eltanexor was combined with EIF4A1 inhibitors (Zotatifin, CR-1-31-B, Rocaglamide A, and C5-desmethyl PatA), and Zotatifin was combined with XPO1 inhibitors (Eltanexor, Selinexor, Verdinexor, and Leptomycin B) across a range of starting concentrations. Excess HSA scores were calculated to quantify synergy across DU145, PC3, LNCaP-95, LNCaP, 22Rv1, VCaP-CR, and VCaP cells. (G-H) Focused 10 × 10 dose-response matrices for the Eltanexor + Zotatifin combination extracted from the combination screen and analyzed using SynergyFinder in (G) LNCaP-95 and (H) VCaP-CR cells. Heatmaps display percent inhibition of cell viability, with corresponding three-dimensional surface plots depicting Highest Single Agent (HSA) synergy scores. (I-K) Validation of Eltanexor + Zotatifin synergy in patient-derived organoid models. Percent inhibition heatmaps and three-dimensional HSA synergy surface plots are shown for (I) LuCaP167-CR, (J) LuCaP136-CR, and (K) Lym1 organoids. Data represent the mean from n=3 replicates. See also Figure S2-4.

Overlaying Excess HSA synergy scores onto the clustered heatmap highlighted synergistic and antagonistic interactions (Figure 2B). Among these, XPO1 inhibitors displayed pronounced synergy with the EIF4A1 inhibitor Zotatifin, which remained highly represented when applying a stringent Excess HSA threshold (Excess HSA < -2000) and visualized as a circos plot (Figure 2B-C). Ranking individual pairs further emphasized this synergy, with Zotatifin emerging as the top partner for both Eltanexor (Figure 2D) and Selinexor (Figure S3A), and reciprocally, Eltanexor and Selinexor ranking first and second among Zotatifin’s most synergistic pairs (Figure 2E).

To assess the robustness of XPO1-EIF4A1 synergy, we performed a targeted combination screen with three objectives: (1) expand testing across a panel of prostate cancer models, including castration-sensitive, castration-resistant, and aggressive variant cell lines (LNCaP, VCaP, 22Rv1, LNCaP-95, VCaP-CR, DU145, and PC3); (2) evaluate synergy across a broad range of concentrations; and (3) determine whether the interaction was mechanistically consistent across structurally distinct inhibitors of the same targets. Notably, correlation analysis of the initial 42 × 42 combination screen revealed only partial concordance between LNCaP-95 and VCaP-CR cells, reflecting cell line-specific differences in single agent sensitivity (Supplementary Figure S3B-D). Therefore, these results, along with single agent dose-response data from Figure 1, S1, and S2, guided the selection of low and high starting concentrations for each drug.

Given the significant synergy observed for Zotatifin + Eltanexor, we selected this pair as our lead combination. In this targeted screen, Eltanexor, as the lead XPO1 inhibitor, was combined with four structurally distinct EIF4A1 inhibitors (Zotatifin, CR-1-31-B, Rocaglamide A, and C5-desmethyl PatA) across a 10 × 10 dose matrix. Conversely, Zotatifin, as the lead EIF4A1 inhibitor, was combined with four XPO1 inhibitors (Eltanexor, Selinexor, Verdinexor, and Leptomycin B) in the same manner. All four possible starting concentration pairings (low-low, low-high, high-low, high-high) were then assessed for each drug combination. Synergy was quantified by Excess HSA, confirming that XPO1-EIF4A1 interactions were reproducible across diverse models, concentrations, and mechanistic analogs (Figure 2F, Table S3). To highlight the clinical relevance of this interaction, we extracted focused matrices for Eltanexor + Zotatifin (Figure 2G-H) and Selinexor + Zotatifin (Figure S3E-F) in LNCaP-95 and VCaP-CR cells with low starting doses (2nM-500nM range). These analyses demonstrated synergy even at reduced concentrations, underscoring potent combination activity with the potential to minimize toxicity.

Importantly, these combinations were manually validated in conventional 96-well plates and orthogonal assays to further confirm the high-throughput screening-identified hits (Figure S4). LNCaP-95, VCaP-CR, and 22Rv1 cells treated with the combinations displayed consistent percent inhibition and Excess HSA synergy scores across 2D cultures (Figure S4A-F). Similarly, 3D spheroids and cell aggregates recapitulated combination-dependent reductions in viability, with single agents producing minimal effects (Figure S4G-J).

Finally, to evaluate translatability in a more physiologically relevant context, we validated Eltanexor + Zotatifin (Figure 2I-K) and Selinexor + Zotatifin (Figure S3G-I) synergy in AR-V7-expressing patient-derived organoid models (LuCaP167-CR, LuCaP136-CR, and Lym1). These organoids recapitulated the potent synergistic interaction observed in cell lines, highlighting the robustness and potential clinical relevance of the XPO1 + EIF4A1-targeting combination.

### Combined inhibition of XPO1 and EIF4A1 induces apoptosis and suppresses proliferation

To explore the mechanism underlying the reduction in cell viability observed with XPO1 + EIF4A1-targeting combinations, we first conducted a high-throughput Caspase-Glo screen combining Zotatifin with XPO1 inhibitors (Eltanexor, Selinexor, Verdinexor, and Leptomycin B) across 10 × 10 dose-response matrices. Caspase activity was measured every 2 hours from 2 to 24 hours post-treatment, allowing quantification of apoptotic responses across both dose and time (Figure 3A, Table S4). This analysis revealed early induction of apoptosis with a maximal response observed at 24 hours, demonstrating that the combination effectively triggers programmed cell death within the first day of treatment. Timing and magnitude of synergy were highlighted by focused views of the Eltanexor + Zotatifin combination; heatmaps and three-dimensional surface plots of HSA scores revealed that apoptosis is strongly synergistic across the combination space (Figure 3B-D).

**Figure 3.**
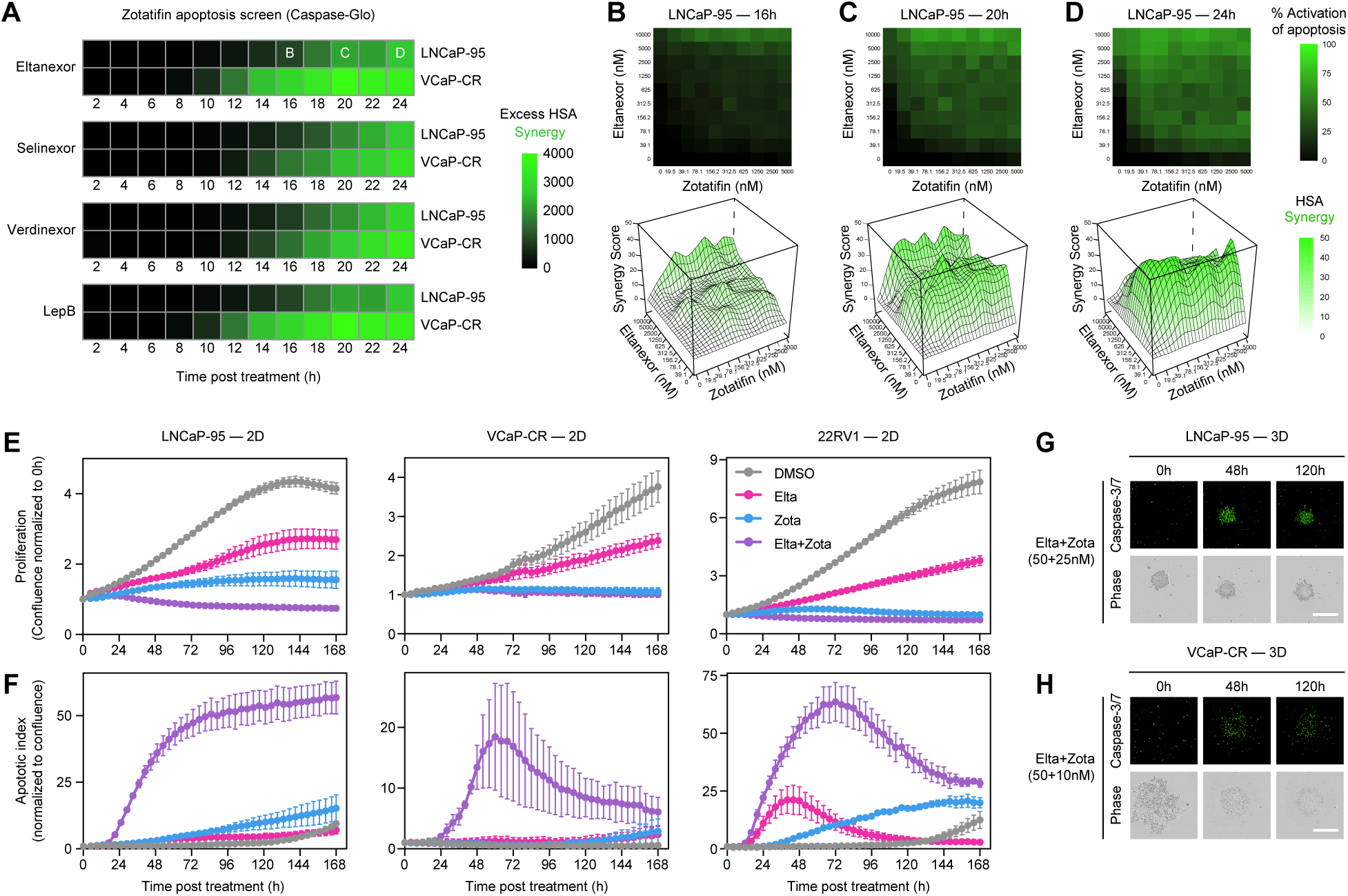
Combined inhibition of XPO1 and EIF4A1 induces apoptosis and suppresses proliferation. (A) Time-course Caspase-Glo screening in which Zotatifin was combined with XPO1 inhibitors (Eltanexor, Selinexor, Verdinexor, and Leptomycin B) in LNCaP-95 and VCaP-CR cells using 10 × 10 dose-response matrices. Caspase activity was measured every 2 hours from 2 to 24 hours post-treatment, and synergy was quantified as Excess Highest Single Agent (Excess HSA). (B-D) Focused analysis of the Zotatifin-Eltanexor combination at (B) 16, (C) 20, and (D) 24 hours in LNCaP-95 cells. Heatmaps display the percent apoptotic response measured by Caspase-Glo, with corresponding three-dimensional surface plots depicting Highest Single Agent (HSA) synergy scores. (E-F) Live-cell imaging (Incucyte) of LNCaP-95, VCaP-CR, and 22Rv1 cells treated with DMSO, single agents, or the Eltanexor + Zotatifin combination. Images were acquired every 4 hours over 7 days. (E) Cell proliferation quantified as percent confluence normalized to 0 h. (F) Apoptotic index quantified using Caspase-3/7 fluorescent dye, normalized to cell confluence. Data represent mean ± SEM from n = 2-3 replicates. (G) Representative Incucyte images of LNCaP-95 cells grown as 3D spheroids and treated with the Eltanexor + Zotatifin combination. Images show Caspase-3/7 fluorescence (green) with corresponding phase-contrast images at 0, 48, and 120 hours. Scale bar, 400 μm. (H) As in (G), representative Incucyte images of VCaP-CR 3D cell aggregates.

To evaluate the long-term impact of XPO1 + EIF4A1 inhibition, we performed live-cell imaging of LNCaP-95, VCaP-CR, and 22RV1 cells treated with low-dose Eltanexor + Zotatifin. Quantification of cell confluence over 7 days revealed that Zotatifin alone reduced proliferation, whereas the combination with Eltanexor further suppressed growth relative to either single agent or DMSO control (Figure 3E). In contrast, concurrent monitoring of Caspase-3/7 activity showed minimal apoptosis for either single agent, with pronounced induction occurring only in the combination treatment (Figure 3F), recapitulating the rapid and synergistic apoptotic effects observed in the short-term Caspase-Glo screen (Figure 3A-D).

We next extended these observations to 3D cultures to better model physiologically relevant tumor architecture. In LNCaP-95 spheroids (Figure 3G) and VCaP-CR aggregates (Figure 3H), the combination of Zotatifin with Eltanexor induced marked structural disruption, with the outer edges of spheroids disintegrating over time. Caspase-3/7 fluorescence confirmed that apoptosis accompanied these morphological changes, reinforcing that dual inhibition is necessary to achieve maximal cytotoxicity (Figure 3G-H). Together, these results demonstrate that XPO1 + EIF4A1-targeting combinations not only suppress proliferation but also trigger synergistic apoptotic responses in both 2D and 3D prostate cancer models.

### Combined inhibition of XPO1 and EIF4A1 drives coordinated suppression of oncogenic programs and cell-cycle signaling

To further interrogate the mechanism underlying the synergistic interaction observed with XPO1 + EIF4A1-targeting combinations, we performed global quantitative proteomic analysis in LNCaP-95 and VCaP-CR cells (Figure S5A). Cells were treated with DMSO, Eltanexor (50 nM), Zotatifin (LNCaP-95: 25 nM; VCaP-CR: 10 nM), or the combination. All treatment conditions were harvested at 24 hours; combination-treated samples were additionally collected at 12 and 48 hours to capture early and late proteomic responses. Protein abundance changes for all conditions were then quantified relative to DMSO controls (Table S5).

Volcano plot analysis revealed that while Eltanexor treatment selectively downregulated XPO1, consistent with its role in nuclear export, Zotatifin impacted protein expression more broadly, reflecting its function as an inhibitor of translation initiation (Figure 4A-B). Notably, Zotatifin treatment led to pronounced downregulation of key oncogenic and metabolic proteins, including AR and SLC2A1 (GLUT1). Combination treatments induced progressively larger changes over time, with significant alterations evident at 12 hours and further increasing at 24 and 48 hours (Figure S5B). AR, SLC2A1, and XPO1 remained suppressed as seen with single agent treatments, while oncogenes such as CDK4 and CCND1 were additionally downregulated in combination treatments (Figure 4A-B, Figure S5B).

**Figure 4.**
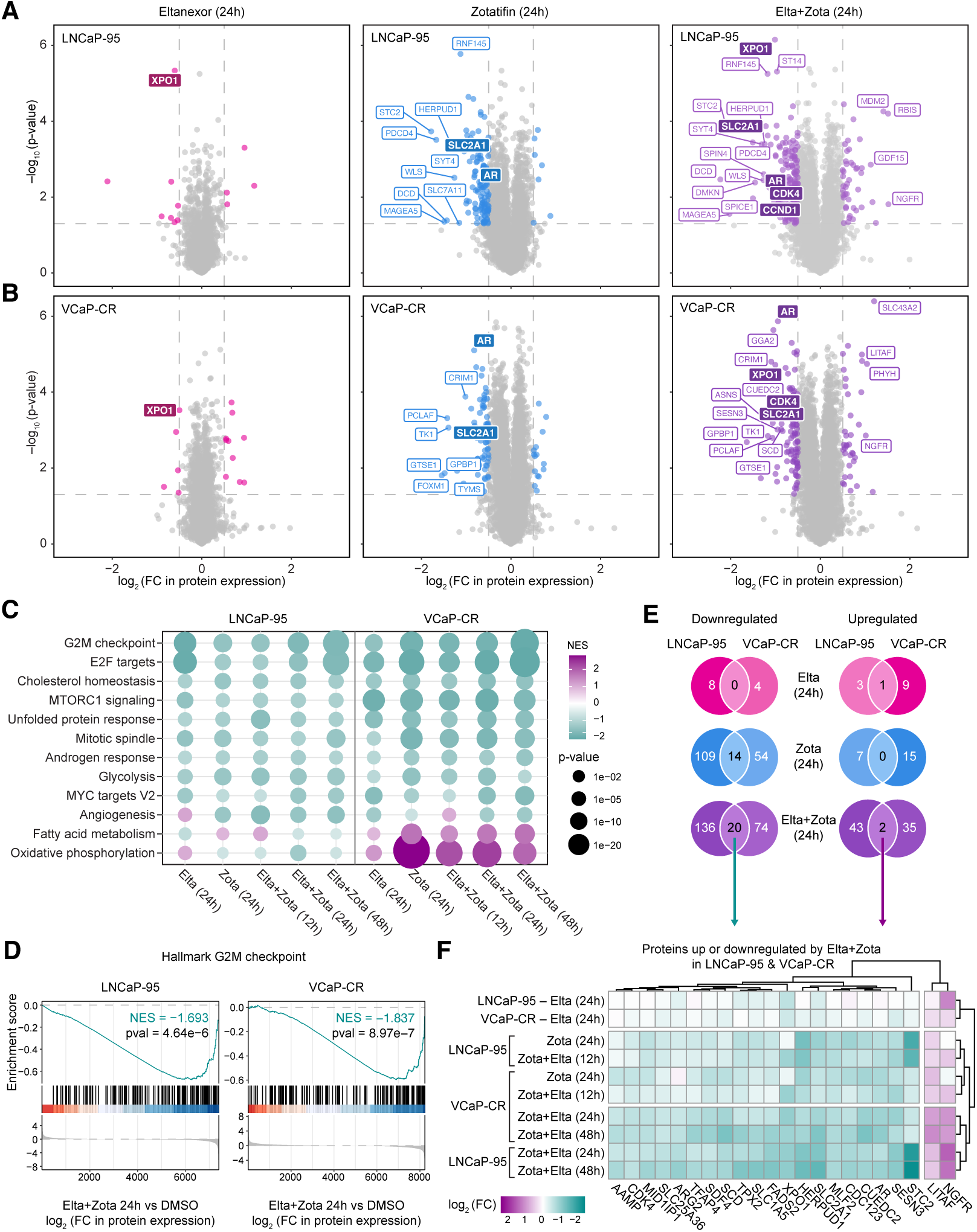
Combined inhibition of XPO1 and EIF4A1 drives coordinated suppression of oncogenic programs and cell-cycle signaling. (A-B) Global quantitative proteomic analysis of (A) LNCaP-95 and (B) VCaP-CR cells treated with Eltanexor (50 nM), Zotatifin (LNCaP-95: 25 nM; VCaP-CR: 10 nM), or the Eltanexor + Zotatifin combination for 24 hours. Data represent the mean from n=3 replicates. Protein abundance changes were assessed relative to DMSO control, and results are displayed as volcano plots depicting log_2_(fold-change) versus -log_10_(p-value). Selected significantly upregulated and downregulated proteins are highlighted. (C) Hallmark pathway enrichment analysis of LNCaP-95 and VCaP-CR proteomic datasets ranked by differential protein abundance for each treatment condition relative to DMSO. Dot plots display normalized enrichment scores for significantly enriched pathways, with dot size indicating statistical significance. (D) Gene set enrichment analysis of the Hallmark G2M checkpoint pathway performed the global proteomics datasets following treatment with the combination for 24 hours, relative to DMSO. Enrichment plots are shown for LNCaP-95 and VCaP-CR cells. (E) Venn diagrams depicting significantly downregulated (log2 fold change < -0.5, p < 0.05) and significantly upregulated proteins (log2 fold change > 0.5, p < 0.05) following Eltanexor, Zotatifin, or combination treatment (24 h) relative to DMSO, highlighting overlapping protein suppression across LNCaP-95 and VCaP-CR cells. (F) Heatmap of proteins commonly regulated by the Eltanexor + Zotatifin combination in both LNCaP-95 and VCaP-CR cells, including 20 overlapping downregulated proteins and 2 overlapping upregulated proteins. Log_2_(fold-changes) are shown across all treatment conditions and cell lines, with hierarchical clustering applied to both proteins and samples to reveal shared regulatory patterns. See also Figure S5.

To characterize coordinated pathway-level effects, we ranked the proteomic datasets relative to DMSO and performed Gene Set Enrichment Analysis (GSEA) using the MSigDB Hallmark pathways. Across treatments and timepoints, common patterns of negatively enriched gene sets emerged, including downregulation of G2M checkpoint, E2F targets, cholesterol homeostasis, mTORC1 signaling, unfolded protein response, mitotic spindle, androgen response, glycolysis, MYC targets, and angiogenesis (Figure 4C-D). Interestingly, VCaP-CR cells displayed concurrent upregulation of oxidative phosphorylation and fatty acid metabolism, consistent with a shift in metabolic programs upon treatment. Furthermore, additional enrichment analyses using MSigDB curated gene sets (C2) and gene ontology biological processes (C5) corroborated suppression of cell cycle, AR, MYC, and E2F signaling programs following XPO1 and EIF4A1 inhibition (Figure S5C-D).

To identify proteins most consistently affected across models, we applied thresholds for significant differential abundance (log_2_ fold-change > 0.5 or < -0.5, p < 0.05) (Table S6). Overlap analysis revealed 20 proteins consistently downregulated and 2 proteins consistently upregulated by the combination in both LNCaP-95 and VCaP-CR cells at 24 hours (Figure 4E-F). Comparison of single agent versus combination-specific effects further identified four proteins downregulated exclusively by the combination across both cell lines: FADS2, SLC25A36, TFAP4, and ARG2 (Figure S5E). These findings indicate that dual inhibition of XPO1 and EIF4A1 not only reinforces the suppression of proteins affected by single agents but also uniquely targets additional oncogenic pathways. Collectively, these proteomic analyses reveal that co-inhibition of XPO1 and EIF4A1 drives coordinated suppression of key oncogenic and cell-cycle signaling programs, providing mechanistic insight into the synergistic reduction in cell viability observed across AR-V7-expressing prostate cancer models.

### Combined XPO1 and EIF4A1 inhibition suppresses AR/AR-V7 signaling and promotes nuclear p53 accumulation

To validate and extend the proteomic findings at a subcellular level, and to directly assess androgen receptor splice variant signaling that could not be resolved in the global proteomics dataset, we performed subcellular fractionation followed by immunoblot analysis in LNCaP-95 and VCaP-CR cells treated with Eltanexor, Zotatifin, or the combination. Separation of cytoplasmic and nuclear fractions allowed direct assessment of protein abundance and localization following treatment (Figure 5A-B).

**Figure 5.**
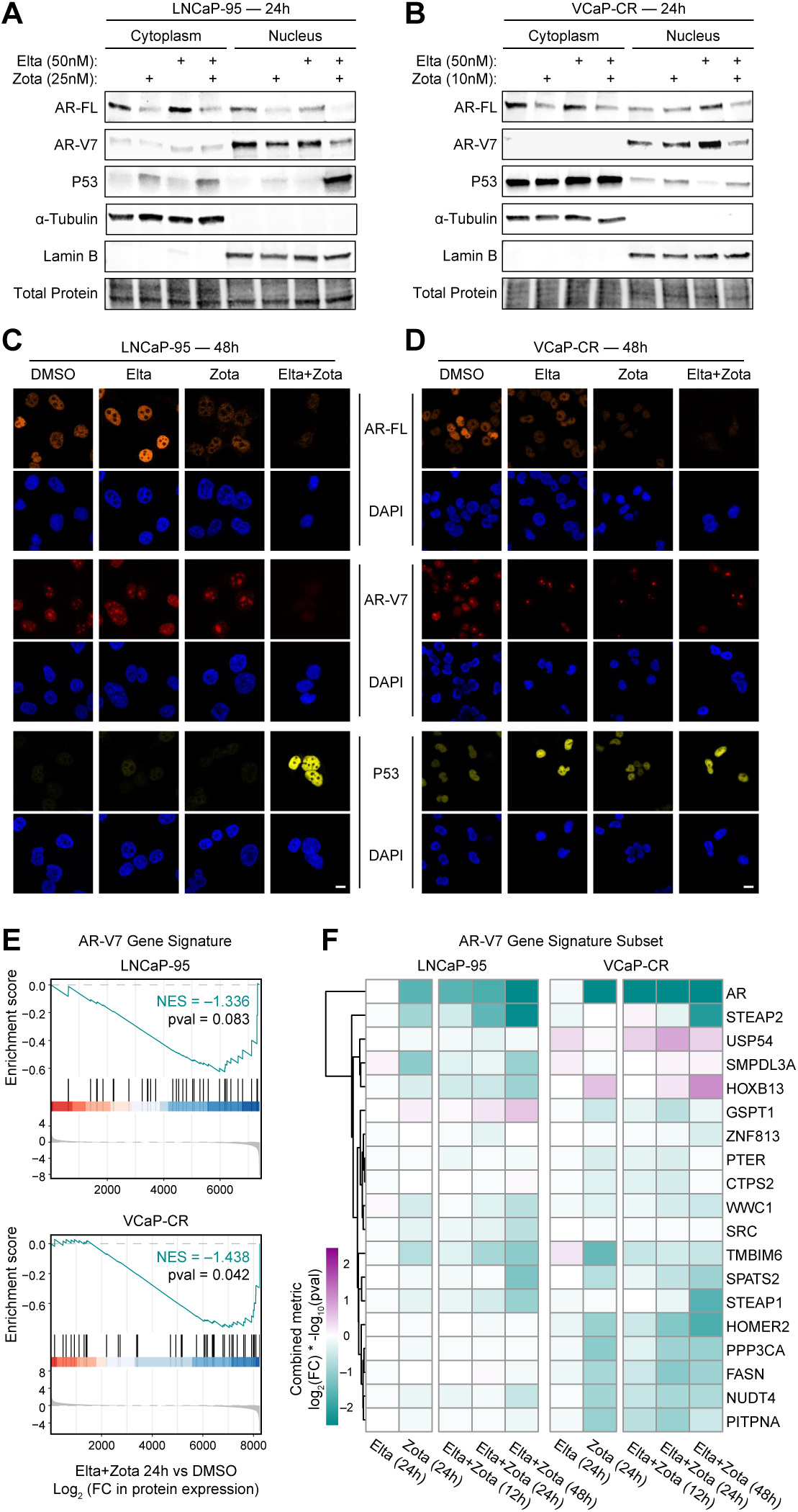
Combined XPO1 and EIF4A1 inhibition suppresses AR/AR-V7 signaling and promotes nuclear p53 accumulation. (A-B) Subcellular fractionation followed by immunoblot analysis of cytoplasmic and nuclear protein extracts from (A) LNCaP-95 and (B) VCaP-CR cells treated with DMSO, Eltanexor, Zotatifin, or the combination for 24 hours. Immunoblots were probed for the indicated proteins, with total protein staining shown as a loading control. (C-D) Immunofluorescence staining of (C) LNCaP-95 and (D) VCaP-CR cells treated with Eltanexor, Zotatifin, or the combination for 48 hours. Cells were stained for the indicated proteins, with DAPI marking nuclei. Representative images were acquired at 60× magnification. Scale bar, 10 μm. (E) Gene set enrichment analysis of the AR-V7 gene signature (Sharp et al. 2019) performed on global proteomics datasets following treatment with the combination for 24 hours, relative to DMSO. Enrichment plots are shown for LNCaP-95 and VCaP-CR cells. (F) Heatmap of AR-V7 signature genes comprising the leading-edge subsets from the Eltanexor + Zotatifin (24 h) versus DMSO enrichment analyses (E) in both LNCaP-95 and VCaP-CR proteomic datasets. Values represent a combined metric of log_2_(fold-change) multiplied by - log_10_(p-value), with hierarchical clustering applied across genes. See also Figure S6.

Consistent with the proteomic results, Zotatifin treatment led to marked downregulation of full-length AR protein, with the most pronounced suppression observed in the combination-treated cells. Importantly, immunoblotting revealed concordant downregulation of AR-V7 protein, which could not be distinguished from full-length AR in the global proteomics analysis. Although AR-V7 suppression was primarily driven by Zotatifin, the effect was greatest in the combination condition, indicating enhanced disruption of androgen receptor signaling with dual XPO1 and EIF4A1 inhibition.

Given prior reports that XPO1 inhibition can promote apoptosis through nuclear retention and activation of p53, we next examined p53 localization and abundance.^14^ Eltanexor alone increased p53 levels, with the combination treatment resulting in a further enhanced accumulation of p53 within the nuclear fraction, particularly in LNCaP-95 cells. These effects were sustained at 48 hours, as confirmed by repeat fractionation experiments (Figure S6A-B).

To visualize these effects and assess any cell-to-cell heterogeneity, we performed immunofluorescence staining in LNCaP-95 and VCaP-CR cells. Consistent with the immunoblot data, combination treatment resulted in a reduction of AR and AR-V7 staining intensity compared to either single agent. In parallel, p53 signal was modestly increased with Eltanexor alone and further enhanced in the combination-treated cells, with prominent nuclear localization (Figure 5C-D; Figure S6C).

Finally, prompted by the observed suppression of AR-V7 protein, we revisited the proteomics datasets to examine transcriptional programs downstream of AR-V7. Gene set enrichment analysis using a previously defined AR-V7 gene signature revealed that AR-V7-regulated genes were enriched among the most downregulated proteins following combination treatment relative to DMSO in both LNCaP-95 and VCaP-CR cells at 24 hours (Figure 5E).^13^ Assessment of this signature across additional time points revealed a consistent pattern of negative enrichment following combination treatment at both early (12 h) and late (48 h) time points (Figure S6D). While AR-V7 pathway enrichment did not reach statistical significance in LNCaP-95 cells at individual time points, the direction and magnitude of enrichment were stable across the time course and mirrored the significant suppression observed in VCaP-CR cells.

To further examine gene-level dynamics underlying this pathway-level effect, we visualized the leading-edge genes from the 24-hour AR-V7 enrichment analysis across all time points in both models (Figure 5F). This analysis illustrated coordinated trends in suppression of AR-V7-regulated genes, with modest cell line- and time-dependent differences in individual gene behavior. Together, these data demonstrate that combined XPO1 and EIF4A1 inhibition disrupts AR and AR-V7 signaling at both the protein and pathway levels while promoting nuclear accumulation of p53, providing mechanistic support for the synergistic anti-tumor effects observed with this combination.

### Eltanexor and Zotatifin combination treatment potently suppresses tumor growth in vivo

To assess the translational potential of combined XPO1 and EIF4A1 inhibition, we evaluated the antitumor activity of Eltanexor and Zotatifin in vivo using both cell line-derived and patient-derived xenograft models of AR-V7-expressing prostate cancer. Dosing regimens were selected based on clinically recommended human doses of each agent, followed by allometric scaling to mouse equivalents and an additional 8-12-fold dose reduction to account for the marked synergy observed in vitro.^15–19^ The final doses used in these studies were substantially below previously established mouse maximum tolerated doses (MTD), corresponding to approximately 30-fold (Eltanexor) and 17.5-fold (Zotatifin) lower exposures.^20–23^

In LNCaP-95 cell-derived xenografts, combination treatment significantly suppressed tumor growth compared to vehicle control and either single agent over three weeks of therapy (Figure 6A). While Eltanexor alone modestly reduced tumor progression, the combination produced a greater antitumor effect, consistent with the synergistic interactions observed in vitro. Similar results were observed in the LuCaP167-CR patient-derived xenograft model, where combination treatment led to pronounced tumor regression over four weeks of treatment (Figure 6B). The superiority of the combination relative to either monotherapy was readily apparent from longitudinal tumor measurements as well as from gross examination of excised tumors at study endpoint (Figure 6E; Figure S7B).

**Figure 6.**
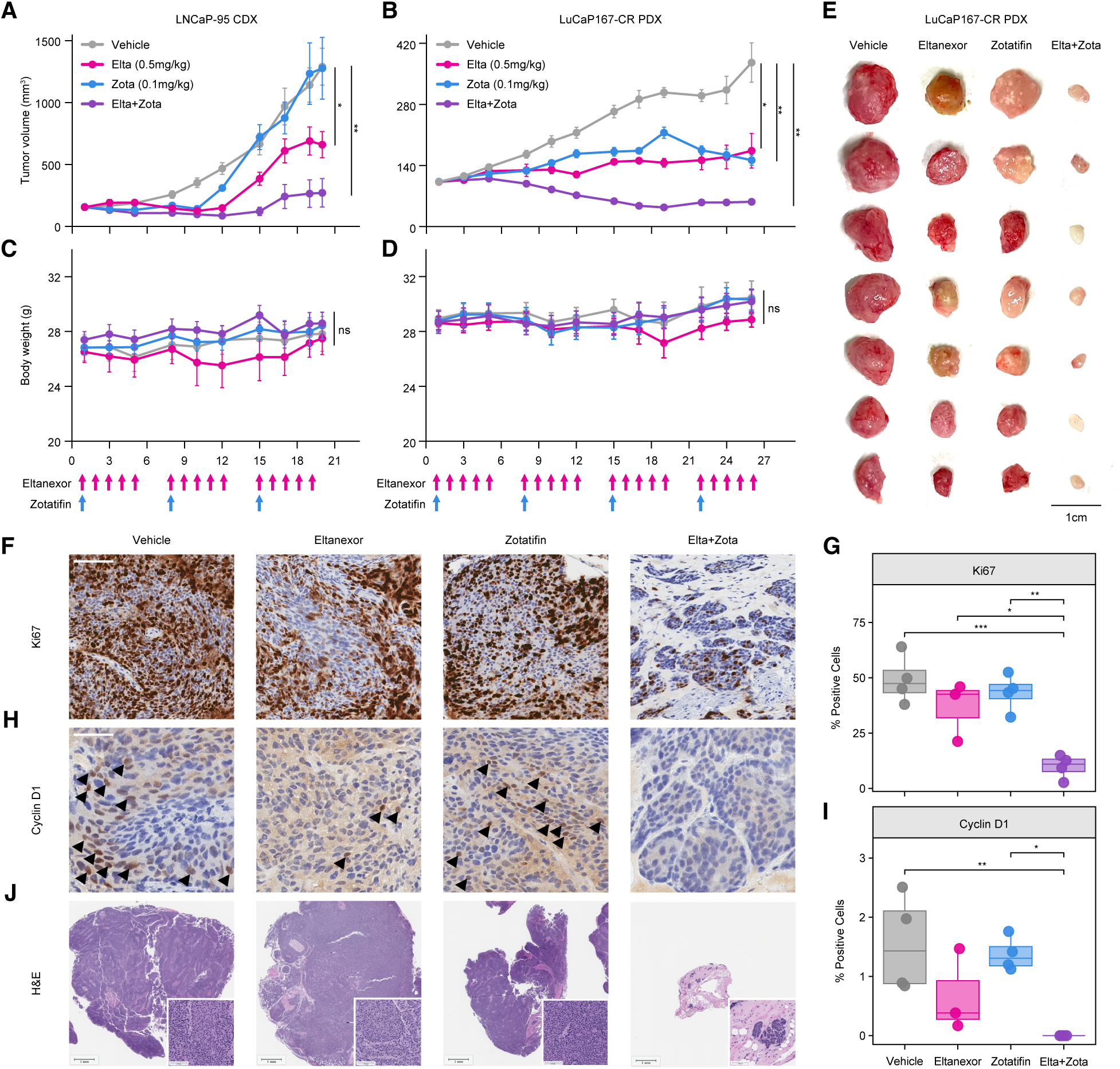
Eltanexor and Zotatifin combination treatment suppresses tumor growth in vivo. (A) Tumor growth curves for LNCaP-95 cell-derived xenograft (CDX) models. Mice were treated with vehicle, Eltanexor (0.5 mg/kg, oral gavage, 5× weekly), Zotatifin (0.1 mg/kg, intravenous, once weekly), or the combination. Tumor volumes were measured over 3 weeks of treatment. Data represent mean ± SEM from n=5 mice per treatment group. Statistical significance was assessed using a mixed-effects model with Sidak’s multiple-comparisons test (*p < 0.05, **p < 0.01). (B) Tumor growth curves for LuCaP167-CR patient-derived xenograft (PDX) models treated with vehicle, Eltanexor, Zotatifin, or the combination for 4 weeks (n=7 mice per treatment group). (C-D) Body weight measurements (g) for mice treated as in (A-B) in (C) LNCaP-95 CDX and (D) LuCaP167-CR PDX studies. (E) Gross images of LuCaP167-CR tumors collected after 4 weeks of treatment. Scale bar, 1 cm. (F) Representative immunohistochemical staining of Ki67 in LuCaP167-CR tumors following treatment. Scale bar, 100 μm. (G) Quantification of Ki67-positive cells across treatment groups (n = 4 tumors per group). Data are shown as box plots with median and distribution; statistical significance was assessed by one-way ANOVA with Tukey’s multiple comparisons test (*p < 0.05, **p < 0.01, ***p < 0.001). (H) Representative immunohistochemical staining of Cyclin D1 in LuCaP167-CR tumors following treatment. Scale bar, 50 μm. (I) Quantification of Cyclin D1-positive nuclei across treatment groups (n = 4 tumors per group). Data are shown as box plots with median and distribution; statistical significance was assessed by one-way ANOVA with Tukey’s multiple comparisons test (*p < 0.05, **p < 0.01, ***p < 0.001). (J) Representative hematoxylin and eosin (H&E) staining of tumors from each treatment group. Insets show higher-magnification views of representative tumor regions. Scale bars, 1 mm (main images) and 100 μm (insets). See also Figure S7.

Importantly, combination treatment was well tolerated in both in vivo models. No significant changes in body weight were observed during treatment in either the CDX or PDX studies (Figure 6C-D), and terminal assessments revealed no differences in body or organ weights between treatment groups (Figure S7A-B). Hematologic and serum chemistry analyses demonstrated no treatment-associated changes outside normal reference ranges in mice receiving single agent or combination therapy, indicating no evidence of overt systemic toxicity (Table S7). These findings indicate that the antitumor efficacy of combined XPO1 and EIF4A1 inhibition can be achieved at doses well below established toxicity thresholds.

To further explore molecular correlates of treatment response in vivo, we performed immunohistochemical analyses of LuCaP167-CR tumors harvested at the end of the treatment period (Figure 6F-J). Combination treatment resulted in a significant reduction in Ki67-positive cells compared to both vehicle control and single-agent groups (Figure 6F-G), consistent with a marked suppression of tumor cell proliferation. Analysis of Cyclin D1 (CCND1) revealed a modest decrease with Eltanexor monotherapy, whereas the combination treatment induced a significant reduction relative to both vehicle control and Zotatifin (Figure 6H-I). These findings are concordant with our proteomic data and support a role for CCND1 downregulation in mediating the enhanced antiproliferative effects of dual XPO1 and EIF4A1 inhibition. Histopathologic evaluation by H&E staining further highlighted the profound impact of combination treatment. Tumors from combination-treated mice were markedly reduced in size and exhibited even further depletion of viable tumor cells, with much of the remaining tissue composed of stromal and adipose elements (Figure 6J).

Together, these in vivo studies demonstrate that low-dose combined inhibition of XPO1 and EIF4A1 produces robust antitumor activity in clinically relevant prostate cancer models while maintaining a favorable tolerability profile, supporting the translational potential of this combination strategy.

## DISCUSSION

In this study, we leveraged a robotically automated high-throughput screening platform to systematically interrogate thousands of mechanistically annotated compounds, enabling the identification of robust single agent activities and high-confidence combinatorial synergies across diverse prostate cancer models. Through this unbiased screening strategy, we identified inhibition of the nuclear export receptor XPO1 as a consistent single agent vulnerability.

XPO1 has emerged as an oncogenic dependency across multiple malignancies, including prostate cancer. As a central mediator of nucleocytoplasmic transport, XPO1 regulates the localization of numerous tumor suppressors, cell-cycle regulators, and components of ribosomal biogenesis; its overexpression promotes tumor cell survival through functional inactivation of nuclear tumor suppressor programs, including attenuation of p53 function and broader dysregulation of nuclear export.^24^ In prostate cancer, XPO1 expression is elevated relative to benign tissue and is associated with aggressive disease features, including higher Gleason score, disease progression, and the development of bone metastases, providing a strong biological rationale for therapeutic targeting.^12,25,26^ These observations motivated the development of selective inhibitors of nuclear export (SINE), which have demonstrated clinical activity in hematologic malignancies. The first-generation SINE agent, Selinexor (KPT-330), was approved by the United States Food and Drug Administration (FDA) for refractory multiple myeloma and showed evidence of antitumor activity in mCRPC patients refractory to anti-androgen therapy (NCT02215161); however, dose-limiting toxicity constrained its therapeutic window, leading to early study termination.^27,28^ Second-generation XPO1 inhibitors, such as Eltanexor (KPT-8602), were subsequently developed to improve safety profiles in solid tumors (NCT02649790) and remain promising candidates for rational combination-based strategies in prostate cancer.^19,29,30^

Importantly, our large combination matrix screen revealed a particularly strong, reproducible, and selective synergy between XPO1 inhibition and Zotatifin, a compound targeting the translation initiation factor EIF4A1, uncovering translational control as a complementary vulnerability. The RNA helicase EIF4A1 is a critical component of translation initiation, facilitating the unwinding of structured 5′ untranslated regions and thereby enabling efficient translation of select proliferation- and survival-associated mRNAs.^31–33^ Pharmacologic inhibition of EIF4A1 using the small-molecule agent Zotatifin (eFT226) has shown favorable tolerability and early signs of biological and clinical activity.^34^ Zotatifin is currently being evaluated in a phase I/II trial in patients with advanced solid tumors (NCT04092673) and in a phase II trial in breast cancer (NCT05101564), where early results suggest antitumor activity in combination with standard-of-care therapy.^15,16^ These developments indicate that targeting translational dependencies may represent a clinically feasible strategy in recalcitrant cancers, including mCRPC.

The convergence of established clinical feasibility, compelling mechanistic rationale, and the pronounced synergy observed in our screen highlighted co-inhibition of XPO1 and EIF4A1 as a therapeutically promising and highly translatable approach for rapidly addressing unmet needs in lethal prostate cancer. Our data demonstrate that co-inhibition of XPO1 and EIF4A1 produces robust synergy that is reproducible in multiple orthogonal assays and across distinct generations of inhibitors targeting each pathway, supporting a drug class-wide effect rather than dependence on a specific compound. This synergy was further validated in advanced model systems, including three-dimensional patient-derived organoids and xenografts, and was biologically conserved across major disease states spanning CSPC, CRPC, and AVPC.

The breadth of this validation is particularly important in advanced prostate cancer, a disease defined by profound molecular heterogeneity and dynamic lineage plasticity. Although TP53 mutation status has been proposed as a predictor of response to XPO1 inhibition alone^35^, XPO1/EIF4A1 co-targeting was consistently efficacious across models with divergent TP53 status (LuCaP167: TP53 wildtype; LuCaP136/Lym1: TP53-/mut; PC3/DU145: TP53 null), indicating that the observed synergy does not depend on p53 reactivation. Further conserved efficacy across androgen-dependent and independent contexts, genetically distinct backgrounds, and patient-derived systems suggests that this combination targets a core regulatory dependency. These consistent responses across diverse preclinical models underscore the translational relevance of XPO1/EIF4A1 co-inhibition for molecularly heterogeneous and heavily pretreated mCRPC populations with pronounced therapeutic resistance.

Perhaps most impressive is the exceptional synergy achieved by the combination at doses substantially lower than the maximum tolerated doses (MTD) reported for Eltanexor or Zotatifin monotherapy, supporting a widened therapeutic window that may mitigate the toxicities previously associated with nuclear export inhibition.^27,28^ To optimize efficacy while minimizing toxicity, we selected combination doses in preclinical xenograft studies in line with the United States Food and Drug Administration’s Project Optimus framework, resulting in exposures 30-fold (Eltanexor) and 17.5-fold (Zotatifin) below the animal MTD.^20–23^ These doses correspond to approximately 8-fold (Eltanexor) and 12-fold (Zotatifin) below the established human single-agent MTD regimens.^15,16,19^

Importantly, robust cooperative pathway suppression was achieved at concentrations where single agents showed minimal activity, indicating that the synergy is not dependent on maximal target inhibition. This provides a mechanistic basis for improved tolerability and is consistent with the broader principle that targeted therapies can achieve efficacy at submaximal doses due to pathway-specific dependencies. Preferential targeting of tumor-specific dependencies, while sparing normal tissues, likely contributes to this effect. Achievement of antitumor efficacy at doses well below those required for single-agent activity is particularly notable, as combination regimens in oncology more commonly exacerbate toxicity rather than improve tolerability. This property may be especially meaningful in mCRPC, where cumulative treatment burden frequently limits further therapeutic escalation.

Mechanistically, dual XPO1 and EIF4A1 inhibition drove concerted suppression of proliferation and induction of apoptosis, accompanied by widespread remodeling of oncogenic protein networks (Figure 7). Global proteomic analyses revealed suppression of key drivers of prostate cancer progression, including AR, CDK4, CCND1, and metabolic regulators such as SLC2A1 (GLUT1), with effects that were amplified by combination treatment. These protein-level changes were mirrored by concordant pathway-level downregulation of cell cycle, androgen response, MYC/E2F, and metabolic programs that support cellular adaptation and survival. Comparison of single agent versus combination-specific effects further identified a discrete subset of proteins uniquely suppressed by dual XPO1 and EIF4A1 inhibition, suggesting that this combination not only reinforces shared oncogenic vulnerabilities but also engages additional metabolic and transcriptional dependencies relevant to cancer biology. These include proteins linked to lipid metabolism (FADS2), mitochondrial transport (SLC25A36), transcriptional regulation (TFAP4), and tumor-immune microenvironment reprogramming (ARG2).^36–41^ Importantly, we also observed marked nuclear accumulation of p53 following combination treatment, linking impaired nuclear export with restored tumor suppressor activity and providing a mechanistic explanation for the enhanced apoptotic response.

**Figure 7.**
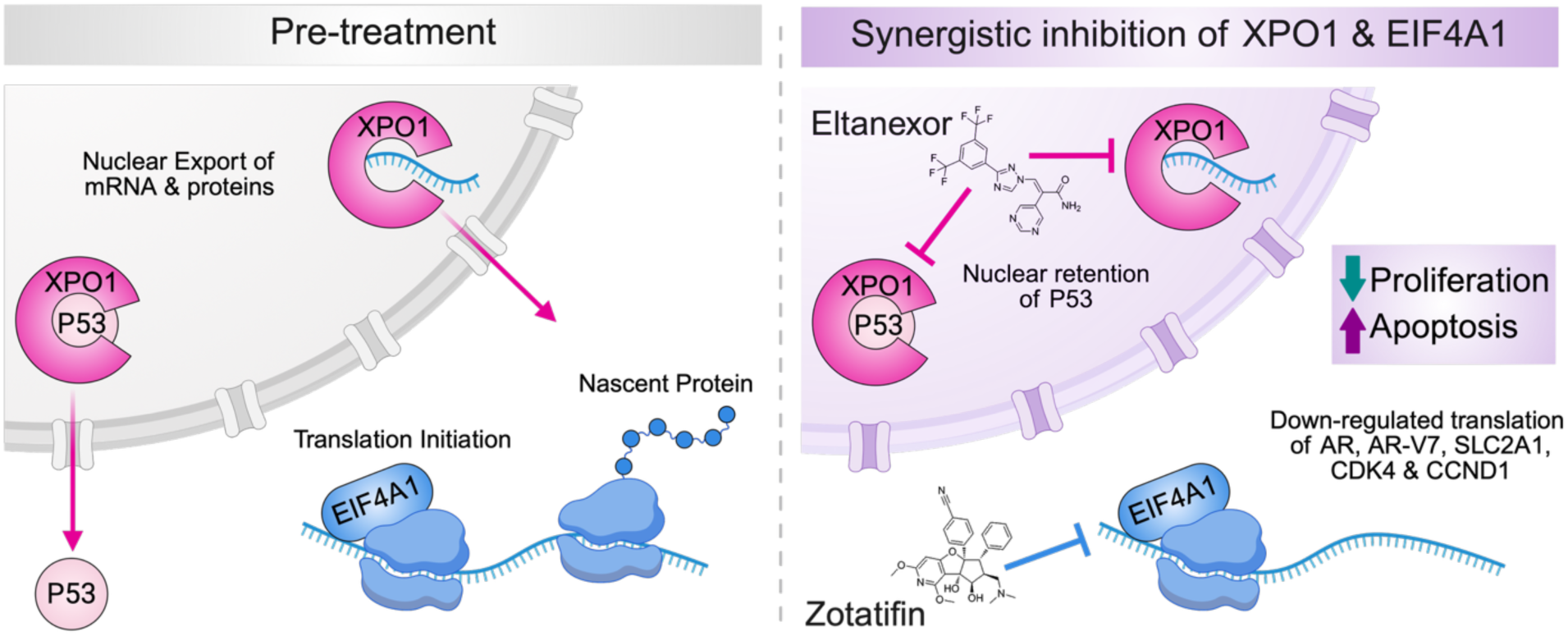
Proposed mechanism underlying synergy between Eltanexor and Zotatifin. Schematic model illustrating the proposed mechanism of synergy between XPO1 and EIF4A1 inhibition. XPO1 inhibition by Eltanexor promotes nuclear retention of p53, leading to the induction of apoptosis, while inhibition of EIF4A1 by Zotatifin suppresses the translation of key oncogenic drivers, including AR, AR-V7, XPO1, SLC2A1, CDK4, MYC, and CCND1 resulting in reduced cell-cycle progression and proliferation. Created with Biorender.com.

While these findings highlight the convergence of nuclear export and translational control as a powerful therapeutic axis, our study also reveals important limitations and directions for future investigation. Our proteomic analyses were performed at the global level and therefore do not fully capture the spatial redistribution of proteins and RNAs that is central to XPO1 function. In addition, targeted marker analyses at the end of the PDX study provide only a single snapshot of tumor burden and may miss transient signaling changes, intermediate target engagement, and the temporal dynamics of drug action. Future studies integrating subcellular fractionation with proteomic and transcriptomic profiling will be essential to define which RNAs and protein complexes are selectively retained in the nucleus following combination treatment. In parallel, longitudinal sampling at earlier timepoints will be important to capture rapid or transient pharmacodynamic effects. Additionally, extending these analyses to models of metastatic dissemination may clarify whether dual XPO1 + EIF4A1 inhibition can delay or prevent disease progression. Given the fundamental roles of nuclear export and translational control in oncogenic signaling, these findings may also have broader relevance in tumor types beyond prostate cancer.

Taken together, our findings identify dual inhibition of XPO1-mediated nuclear export and EIF4A1-dependent translation as a potent, mechanistically coherent, and clinically actionable strategy for mCRPC. By targeting complementary vulnerabilities, this combination elicits robust and reproducible synergistic antitumor effects across diverse prostate cancer models while enabling dose reductions that may mitigate toxicity and constrain adaptive resistance. More broadly, these results demonstrate that coordinated disruption of multiple oncogenic processes can overcome the heterogeneity and treatment-refractory nature of advanced prostate cancer, and highlight the value of unbiased, large-scale combination screening for uncovering therapeutically relevant interactions. Collectively, this work provides a strong rationale for clinical evaluation of combined XPO1 and EIF4A inhibition and has motivated our advancement of this strategy into early-phase clinical trials for patients with lethal prostate cancer.

## METHODS

### Cell lines

The human prostate cancer cell lines 22Rv1 (RRID: CVCL_1045), DU145 (RRID: CVCL_0105), PC3 (RRID: CVCL_0035), VCaP (RRID: CVCL_2235), and LNCaP (RRID: CVCL_1379) were obtained from the American Type Culture Collection (ATCC). LNCaP-95 (RRID: CVCL_ZC87) and VCaP-CR (RRID: CVCL_A0BX) cells were kindly provided by Dr. Jun Luo (Johns Hopkins Biobank). LNCaP-abl (RRID: CVCL_4793) cells were kindly provided by Dr. Zoran Culig (Medical University of Innsbruck). Unless otherwise specified, cell culture reagents were purchased from Thermo Fisher Scientific (Gibco). 22Rv1 and LNCaP parental cells were cultured in RPMI-1640 with L-glutamine supplemented with 10% Premium Select fetal bovine serum (FBS; R&D Systems) and 1% penicillin/streptomycin (P/S). LNCaP-95 and LNCaP-Abl cells were maintained in phenol red-free RPMI-1640 supplemented with 10% charcoal stripped FBS (CSS-FBS; R&D Systems) and 1% P/S. VCaP parental cells were cultured in DMEM containing 4.5 g/L D-glucose and L-glutamine supplemented with 10% Premium Select FBS and 1% P/S. VCaP-CR cells were maintained in phenol red-free RPMI-1640 with L-glutamine supplemented with 10% charcoal dextran-stripped FBS and B-27 supplement (Gibco). PC3 cells were cultured in F-12K Nutrient Mixture supplemented with 10% Premium Select FBS and 1% P/S. DU145 cells were maintained in EMEM (ATCC) supplemented with 10% Premium Select FBS and 1% P/S. LNCaP-95 and VCaP-CR cells were grown on Cell+ culture flasks (Sarstedt), while other cell lines were maintained on standard tissue culture-treated plasticware. All cells were maintained at 37°C in a humidified incubator with 5% CO₂. Cell line authentication was performed by short tandem repeat profiling (ATCC/LabCorp). Mycoplasma testing was conducted monthly using the MycoAlert Mycoplasma Detection Kit (Lonza). Cells were used for no longer than three months after thawing.

### Animal experiments

Male immunodeficient NOD/SCID gamma (NSG) mice at 6-8 weeks were used for preclinical trials. NSG mice were anesthetized with ketamine/xylazine (100/10 mg/kg, i.p.). Both testes were moved to the scrotal sacs by gently applying pressure to the abdomen. An 8-10 mm incision through the skin was made along the midline of the scrotal sac. Another incision was made into the midline wall between the testicular sacs under the covering membranes. The testis, the vas deferens, and the epididymal fat pad were pulled carefully out through the incision. The blood vessels supplying the testis were clamped with a hemostat and the testis was dissected away. The vas deferens and the fat pad were cauterized and placed back into the scrotal sac. This procedure was repeated for the other testis. The incision through the skin was closed using wound clips.

LNCaP-95 cells and LuCaP167-CR (3 × 10^6^) were mixed with Matrigel (75 μl of 1× PBS plus 75 μl of Matrigel (Corning)) and injected subcutaneously into the right flank of 8-week-old castrated NSG mice. After xenografts reached a size of ∼100 mm^3^, animals were randomized based on tumor volume and body weight into one of four treatment groups (n = 5-7 per group) including vehicle, Zotatifin (0.1 mg/kg, IV, once per week; formulation: 5% dextrose), Eltanexor (0.5 mg/kg, PO, daily for 5 days on/2 days off; formulation: 1% Tween-80, 0.5% methylcellulose), and the combination of Zotatifin plus Eltanexor. Mice were treated for 3 (LNCaP-95) or 4 (LuCaP167-CR) weeks, with weight and tumor volumes measured three times weekly, using the formula V=(L*W^2^)/2 (length corresponding to longer dimension of tumor). Following treatment conclusion, the mice were euthanized, and tumors were harvested to obtain final weight and volume (using L*W*H*(π/6)) measurements.

At study termination, blood samples were collected via cardiac puncture under terminal anesthesia and processed for hematological and serum chemistry analyses. Whole blood was collected into EDTA-coated tubes for complete blood count (CBC) analysis, including measurements of white blood cells, red blood cells, hemoglobin, hematocrit, platelet counts, and differential leukocyte populations. CBC analysis was performed using an automated hematology analyzer following standard operating procedures. For serum chemistry analysis, blood was collected into serum separator tubes and allowed to clot at room temperature prior to centrifugation. Serum was isolated and analyzed for markers of hepatic and renal function, including alanine aminotransferase (ALT), aspartate aminotransferase (AST), alkaline phosphatase (ALP), total bilirubin, blood urea nitrogen (BUN), and creatinine. Additional metabolic parameters, including glucose, total protein, and albumin, were measured as indicated. All biochemical analyses were performed using a clinical chemistry analyzer according to the manufacturer’s protocols.

All animals were housed in a pathogen-free facility of the National Cancer Institute, which is accredited by the Association for Assessment and Accreditation of Laboratory Animal Care (AAALAC) International and follows the Public Health Service (PHS) Policy for the Care and Use of Laboratory Animals. All animal procedures were performed in accordance with the Guide for the Care and Use of Laboratory Animals. The study protocol was approved by the NCI Animal Care and Use Committee (ACUC).

### Single agent quantitative high-throughput screening

Single agent qHTS was performed in 1,536-well white tissue culture-treated plates. LNCaP-95 or VCaP-CR cells were dispensed in 5 µl of medium per well using a Multidrop automated liquid handling system at a density of 500 cells per well. Compounds from the MIPE 5.0 library were transferred into assay plates using a 1,536-pin tool (23 nl per well). Each compound was tested across an 11-point concentration series using a fixed 1:3 serial dilution ranging from 45 µM to 0.8 nM. Negative controls included DMSO and empty wells, while Bortezomib served as a standard cytotoxic positive control. Plates were incubated under standard culture conditions for 48 h (LNCaP-95 and 22Rv1) or 72 h (VCaP-CR) with stainless steel gasketed lids to minimize evaporation. At the assay endpoint, CellTiter-Glo reagent (Promega) was added, and plates were incubated at room temperature to allow complete cell lysis. Luminescence was measured using a ViewLux plate reader (PerkinElmer) with a 2-second exposure per plate. Single agent dose-response curves were normalized to DMSO and empty well controls. The area under the curve (AUC) for each compound was Z-transformed (Z-AUC) to generate activity scores. Z-AUC values were averaged across LNCaP-95 and VCaP-CR cells and ranked from low to high activity, with higher activity corresponding to lower Z-AUC values. Compounds were visualized using pheatmap (v1.0.13) or ggplot2 (v3.5.2), with each compound labeled according to its annotated mechanistic target (suffix “i” denotes inhibitor). Gene set enrichment analysis (GSEA) was conducted using gseaplot2 on a custom gene set comprising the five XPO1 inhibitors within the MIPE 5.0 library, applied to the preranked Z-AUC file.^42^ Individual dose-response curves were further analyzed in additional cell lines using the R package DRC (v3.0.1) with the drm function, fitting four-parameter logistic (4PL) curves per cell line.^43^

### High-throughput combination drug screening

Pairwise drug combination screens were conducted under the same culture and assay conditions as single agent qHTS, with modifications for matrix-based testing. Forty-two compounds were selected based on single agent activity and mechanistic interest, and all possible pairwise interactions (n = 861) were evaluated using 10 × 10 dose matrices (Table S2). Compounds were acoustically dispensed into 1,536-well plates using an Echo acoustic liquid handler (10 nl per well), generating nine-point concentration series for each drug with 1:2 serial dilutions. Plates were incubated for 48 h (LNCaP-95) or 72 h (VCaP-CR) with stainless steel gasketed lids to prevent evaporation. Cell viability was assessed using CellTiter-Glo (Promega), with 3 µl added per well and plates incubated at room temperature for 15 min prior to luminescence measurement on a ViewLux imager (PerkinElmer, 2-second exposure per plate). Single agent dose-response data were used to guide the selection of starting concentrations for these combinations. Synergy and antagonism were quantified using the Excess Highest Single Agent (Excess HSA) metric as previously described.^10^ Excess HSA values were averaged across LNCaP-95 and VCaP-CR cells for downstream analyses. Unsupervised hierarchical clustering of compounds based on Pearson correlation of Excess HSA profiles was performed using the cor function in R, and heatmaps were visualized with pheatmap (v1.0.13). Excess HSA synergy scores were overlaid on the clustered heatmap to highlight interactions. The most significant combinations (Excess HSA < -2000) were then visualized using circlize (v0.4.16) to generate chord diagrams.^44^ For individual combinations of interest, response matrices were analyzed with SynergyFinder+ (v3.18.0) in RStudio.^45^ Two-dimensional heatmaps of percent inhibition were generated using plot2drugheatmap (plot_value = “response”), and three-dimensional synergy surface plots were generated using plot2drugsurface (plot_value = “HSA_synergy”), referencing the highest single agent model.

#### Targeted combination viability screening

To validate lead combinations and explore mechanistic consistency, selected drug pairs were tested across additional prostate cancer models, including castration-sensitive, castration-resistant, and AR-null cell lines (LNCaP, VCaP, 22Rv1, LNCaP-95, VCaP-CR, DU145, and PC3). Four starting concentration pairings (low-low, low-high, high-low, high-high) were evaluated in 10 × 10 matrices (Table S3). Dose-response data were analyzed and visualized using the same workflow as above.

#### Targeted combination apoptosis screening

Apoptotic activation was assessed for a subset of targeted combinations (Zotatifin paired with Eltanexor, Selinexor, Verdinexor, or Leptomycin B) using Caspase-Glo 3/7 (Promega) following the same plating and treatment procedures as for pairwise drug viability measurements. To capture kinetic changes, and because the Caspase-Glo 3/7 assay is lysis-based, multiple identical plates were prepared, and luminescence readings were collected every 2 h over the duration of the experiment (10-second exposure per plate, ViewLux imager). All other assay conditions and analysis methods were identical to those described for viability measurements.

### Manual cell viability assays

For 2D viability assays, cells were seeded on clear-bottom polystyrene or amine-coated black 96-well plates at cell line-specific densities: LNCaP-95, 15,000 cells/well; 22Rv1, 15,000 cells/well; and VCaP-CR, 20,000 cells/well, in a total volume of 100 µl per well. 3D spheroids were generated in 96-well U-bottom Ultra-Low Attachment Plates (Corning) at the following seeding densities: LNCaP-95, 600 cells/well; 22Rv1, 2,000 cells/well; VCaP-CR, 1,250 cells/well. After 24 hours, the medium was replaced with treatment medium containing either DMSO control or test compounds, administered alone or in combination. Eltanexor (S8397) and Selinexor (S7252) were obtained from SelleckChem and Zotatifin (HY-112163) was obtained from MedChemExpress. Cells were incubated for 48 h (LNCaP-95 and 22Rv1) or 72 h (VCaP-CR), unless otherwise specified. Cell viability was assessed using the 2D or 3D CellTiter-Glo assay (Promega) according to the manufacturer’s instructions. Luminescence was measured with an integration time of 1000 ms using a SpectraMax iD3 plate reader (Molecular Devices). Experiments were performed in 3 biological replicates, with at least 3 technical replicates per condition. Luminescence values were normalized to the DMSO-treated control group. Single agent dose-response curves for 2D experiments were generated in GraphPad Prism using the nonlinear regression fit, log(inhibitor) vs. response (three-parameter). For 3D spheroid experiments, viability data were presented as bar graphs, and statistical significance was assessed using ordinary one-way ANOVA with multiple comparisons, testing DMSO- or single agent-treated cells against combination-treated cells. Manually conducted combination experiments were analyzed using SynergyFinder+, following the workflow described above for matrix-based screens.

### Organoid drug screening

Patient-derived xenograft (PDX) models LuCaP167 and LuCaP136 were obtained from the NCI Patient-Derived Models Repository (PDMR; models K14711-072-R and K31318-253-M, respectively). The original LuCaP167 and LuCaP136 models were developed at the University of Washington.^46,47^ The Lym-1 model used in this study was generated by the National Cancer Institute and kindly provided by Dr. Adam Sowolsky.^48^ Castration-resistant derivatives (LuCaP167-CR and LuCaP136-CR) were established as previously described.^47,49^ Briefly, LuCaP167 and LuCaP136 tumors were implanted into 6-7-week-old NOD/SCID gamma (NSG) mice, followed by surgical castration through scrotal orchiectomy. Resulting tumors were harvested and serially passaged in pre-castrated NSG mice to generate stable castration-resistant PDX lines. For organoid drug screening, tumor tissues were processed as described previously.^46,49^ Tumors were finely minced and enzymatically dissociated in Advanced DMEM using the GentleMACS Tumor Dissociation Kit (Miltenyi Biotec) according to the manufacturer’s instructions. Single-cell suspensions were filtered through 100 μm cell strainers, subjected to red blood cell lysis (Lonza), washed in DMEM supplemented with 10% fetal bovine serum, and subsequently filtered through 70 μm strainers. Cells were centrifuged, counted, and cryopreserved (5 × 10⁶ cells in 90% FBS and 10% DMSO) or immediately used for drug screening. Cells were embedded in 70% growth factor-reduced, phenol red-free Matrigel (Corning) and seeded into 384-well plates at a density of 2,000 cells per well in a total volume of 20 μl. Prostate cancer organoids were cultured in Advanced DMEM/F12 supplemented with 1% penicillin/streptomycin, 1% GlutaMAX, 10 mM HEPES, 50× B-27, 5% R-spondin-conditioned medium, 10% Noggin-conditioned medium, 5 ng/mL EGF, 100 ng/mL FGF-10, 1 ng/mL FGF-2, 10 μM Y-27632, 10 mM nicotinamide, 1 μM prostaglandin E2, and 1 nM dihydrotestosterone. Twenty-four hours after seeding, organoids were treated with Zotatifin and Eltanexor across a concentration range of 1-1000 nM using a 7 × 7 dose matrix dispensed with a Tecan D300e Digital Dispenser. Drug-containing media was refreshed every 3 days. After 10 days of treatment, cell viability was assessed using CellTiter-Glo 3D (Promega) according to the manufacturer’s instructions, and luminescence was measured using an Infinite M200 PRO microplate reader (Tecan). Luminescence values were normalized to DMSO-treated controls and for individual combinations response matrices were analyzed with SynergyFinder+ as described above. All experiments were performed with three independent biological replicates, each containing six technical replicates.

### Live cell imaging

Live-cell imaging was performed using the Incucyte S3 system (Sartorius) to monitor cell proliferation and apoptosis following treatment. For 2D cultures, cells were seeded in 96-well plates at cell line-specific densities: LNCaP-95 and 22Rv1, 3,000 cells/well; VCaP-CR, 4,500 cells/well, in a final volume of 100 µl per well. Incucyte Caspase 3/7 green dye (Sartorius) was added at a 1:2000 dilution prior to plating. Cells were allowed to attach for 24 hours (LNCaP-95 and 22Rv1) or 48 hours (VCaP-CR) prior to treatment with DMSO control or test compounds. Proliferation was quantified using phase contrast with classic confluence-based segmentation, while apoptosis was quantified using top-hat segmentation of Caspase 3/7 fluorescent signal (radius = 10 µm; threshold = 1.4; edge split off). Apoptotic indices were calculated as the number of Caspase 3/7 green-positive cells (>20 µm²) divided by percent confluence (segmentation adjustment = 0.4; filter ≥100 µm²). Images were acquired at regular intervals, and data were analyzed using the Incucyte Basic Analyzer. Confluence and apoptotic signals were normalized to baseline (0-hour) measurements, and final plots were made in GraphPad Prism. For 3D spheroid experiments, cells were seeded in 96-well U-bottom Ultra-Low Attachment Plates (Corning) at the following densities: LNCaP-95, 750 cells/well; 22Rv1, 2,000 cells/well; and VCaP-CR, 1,250 cells/well, in 100 µl of medium. During seeding, Caspase 3/7 dye was added at 1:2000, and spheroids were allowed to form for 24 hours (LNCaP-95 and 22Rv1) or 48 hours (VCaP-CR) hours prior to treatment. Medium was partially replaced with compound-containing media to maintain spheroid integrity. Images were acquired at regular intervals using the Incucyte S3 system and extracted visualization. Separately, select spheroid images for supplemental figures (Figure S2D and Figure S4I-J) were acquired manually using a Nikon Eclipse TE2000-U microscope with NIS-Elements software at 4× magnification (1.6125 µm/pixel). Spheroid diameters were measured using ImageJ (RRID: SCR_003070) to ensure sizes between 200 and 500 µm prior to treatment. All experiments were performed with 2-3 independent biological replicates, each containing a minimum of 3 technical replicates.

### Protein harvesting and subcellular fractionation

8 million cells were seeded per T-75 Cell+ plate (Sarstedt) and allowed to incubate overnight. The cells were treated with DMSO or compounds for 24 or 48 hours. Total protein extracts were prepared by washing cells with ice-cold PBS, then lysing with RIPA lysis buffer (Sigma) supplemented with complete protease inhibitor cocktail (Nacalai). Samples were left on ice for 10 minutes, briefly vortexed, and returned to ice for another 10 minutes. Lysed cells were then centrifuged at 13,000 × g at 4°C for 15 minutes. Subcellular Fractionation of proteins was performed according to the manufacturer’s guidelines to isolate cytoplasmic and nuclear proteins with the NE-PER Nuclear and Cytoplasmic Extraction Reagents kit (Thermo Scientific). Additionally, the nuclear pellet was washed twice with ice-cold CERI to increase fraction purity.

### Western blot analysis

Protein concentration was quantified by Pierce Bicinchoninic Acid (BCA) assay (Thermo Fisher Scientific). Samples were loaded onto 10% SDS-PAGE Mini-PROTEAN TGX™ Precast Gels (Bio-Rad) and separated by electrophoresis at 180 V for 30 minutes. Proteins were then transferred to 0.2 μm nitrocellulose membranes using the Mini Trans-Blot® Turbo™ semi-dry transfer system (Bio-Rad). Before blocking, a total protein stain was performed with the LICOR Revert 700 Total Protein Stain Kit; (Single-Color Western Blot 800 nm target only). After total protein staining, membranes were blocked with 5% nonfat dry milk in TBS with 0.1% Tween (TBST) for 60 minutes, then incubated overnight at 4°C with primary antibodies. Primary antibodies were made in 5% Bovine Serum Albumin (Sigma Aldrich) in TBST according to the following indicated dilutions: AR (SC-7305; 1:200; Lot B0623; Santa Cruz Biotechnology), AR-V7 (RevMAb RM7; 1:1000; Lot U-03-04124; RevMAb Biosciences), p53 (SC-126; 1:1000; Lot B2824; Santa Cruz Biotechnology), XPO1/CRM1 (SC-74454; 1:1000; Lot H0421; Santa Cruz Biotechnology), GAPDH (SC-47724; 1:1000; Lot G1522; Santa Cruz Biotechnology), eIF4A1 (2409S; 1:1000; Lot 2; Cell Signaling Technology), c-MYC (940S; 1:1000; Lot 12; Cell Signaling Technology), Lamin B1 (SC-126; 1:1000; Lot 2901060; Santa Cruz Biotechnology), alpha-tubulin (2144S; 1:2000; Lot 3; Cell Signaling Technology). Membranes were then washed five times with TBST and incubated for one hour at room temperature with LICOR secondary antibodies. Secondary antibodies were made in 5% Milk in TBST using Goat Anti-Mouse IRDye 680RD (1:20000, Licor 925-68070) and Goat Anti-Rabbit IRDye 800CW (1:20000, Licor 925-32211). After three washes with TBST, blots were imaged using the LICOR Odyssey Fc. To enable re-probing, membranes were stripped for 20 minutes with NewBlot IR Stripping Buffer (LICOR) diluted 5x in ultrapure water. Next, the membrane was washed with TBST for three 5-minute washes and 30 minutes of 5% BSA blocking before repeating the procedure above.

### Mass spectrometry

#### Protein digestion and TMT labeling

Cells were collected at the indicated treatment time points for global proteomic analysis. During collection, plates were maintained on ice, washed with ice-cold PBS to remove residual media, and cells were scraped, pelleted by centrifugation, and snap-frozen in liquid nitrogen prior to storage at −80°C. Cell pellets were lysed in EasyPrep Lysis buffer (Thermo Fisher) following the manufacturer’s instructions. Lysates were clarified by centrifugation, and protein concentrations were determined using the BCA protein estimation kit (Thermo Fisher). Fifteen micrograms of protein lysate were subjected to reduction, alkylation, and overnight digestion at 37°C following addition of trypsin at a 1:50 ratio (Promega). For TMT labeling, 100 μg of TMTpro reagent (Thermo Fisher) prepared in 100% acetonitrile (ACN) was added to each sample. Samples were incubated for 1 hour at room temperature with intermittent mixing, after which the reaction was quenched by addition of 50 μl of 5% hydroxylamine and 20% formic acid. Peptides corresponding to each condition were pooled, and clean-up was performed using the proprietary peptide clean-up columns included in the EasyPEP Mini MS Sample Prep kit (Thermo Fisher).

#### High pH reverse phase fractionation

Peptide fractionation was performed as a first-dimension separation using a Waters Acquity UPLC system equipped with a fluorescence detector (Waters) and a 150 mm × 3.0 mm XBridge Peptide BEH™ C18 column (2.5 μm; Waters), operated at a flow rate of 0.35 ml/min. Dried peptide samples were reconstituted in 100 μl of mobile phase A (10 mM ammonium formate, pH 9.4), while mobile phase B consisted of 10 mM ammonium formate in 90% acetonitrile (pH 9.4). The column was equilibrated with mobile phase A for 5 minutes prior to gradient elution (10–50% B from 5–60 minutes, followed by 50–75% B from 60–70 minutes). Fractions were collected at 1-minute intervals, yielding 60 fractions, which were subsequently pooled into 24 fractions. Pooled fractions were dried by vacuum centrifugation and stored at −80°C until mass spectrometry analysis.

#### Mass spectrometry acquisition and analysis

Dried peptide fractions were reconstituted in 0.1% TFA and analyzed by nanoflow liquid chromatography using a Thermo Easy nLC 1200 system (Thermo Scientific) coupled to an Orbitrap Lumos mass spectrometer (Thermo Scientific). Peptides were separated using a low-pH gradient (5–50% ACN over 180 minutes) in a mobile phase containing 0.1% formic acid at a flow rate of 300 nL/min.

Full MS scans were acquired in the Orbitrap at a resolution of 120,000, with an ion accumulation target of 4e5 and a maximum injection time of 50 ms across a mass range of 400–1600 m/z. Precursor ions with charge states between 2 and 5 were selected for MS2 analysis. MS/MS acquisition was performed using a 3-second cycle time and a quadrupole isolation window of 0.7 m/z. Fragment ions were analyzed in the Orbitrap at a resolution of 50,000 with a normalized AGC target of 250, maximum injection time set to Auto, and normalized collision energy of 38.

MS/MS spectra were searched against a human UniProt protein database using SEQUEST HT with Percolator validation implemented in Proteome Discoverer 2.4 (Thermo Scientific). Precursor mass tolerance was set to 10 ppm and fragment ion tolerance to 0.02 Da. Methionine oxidation was included as a dynamic modification, while carbamidomethylation of cysteine residues and TMT16plex labeling (304.2071 Da) on lysine residues and peptide N-termini were specified as static modifications. Trypsin was defined as the proteolytic enzyme with up to two missed cleavages permitted.

A reverse sequence decoy database strategy was used to control false discovery rates, and peptide identifications were validated using Percolator. Only peptides with less than 50% co-isolation interference were retained for quantification. Reporter ion intensities were corrected for isotopic impurities based on manufacturer specifications, and protein abundances were calculated by summing reporter ion intensities across all corresponding peptides. Reporter intensities were normalized across all channels to account for equal peptide loading.

### Proteomics data analysis

For each treatment condition, log2 fold-change (log2FC) values relative to DMSO controls were calculated using the mean of three independent biological replicates. Corresponding p-values were derived from replicate-level comparisons as described above. Differentially abundant proteins were defined using thresholds of |log2FC| > 0.5 and p < 0.05. Volcano plots and overlap analyses were generated using ggplot2 (v3.5.2), ggvenn (v0.1.10), and ggVennDiagram (v1.5.4) in R (v4.5.1). For pathway-level analyses, proteins were ranked using a composite metric defined as log2FC × (-log10 p-value), thereby prioritizing proteins exhibiting both magnitude and statistical significance of differential abundance. Gene set enrichment analysis (GSEA) was performed using preranked protein lists and the fgsea package (v1.34.2), implementing the fast multilevel algorithm.^50^ Gene sets were obtained from the Molecular Signatures Database (MSigDB) using msigdbr (v25.1.1), including Hallmark (H), curated (C2; KEGG, Reactome, PID, BioCarta, WikiPathways), and Gene Ontology Biological Process (C5) collections.^42,51^ Normalized enrichment scores (NES), nominal p-values, and Benjamini-Hochberg FDR-adjusted p-values were calculated for each gene set. Statistical significance was defined as FDR-adjusted p-value (padj) < 0.05. For visualization across multiple treatment conditions and cell lines, dot plots display NES (color) and -log10 nominal p-value (size) without filtering to enable direct comparison of pathway activity patterns. Individual enrichment plots were generated using gseaplot2 from enrichplot (v1.28.4) in conjunction with clusterProfiler (v4.16.0).^52^ To assess androgen receptor splice variant signaling, a custom AR-V7 transcriptional gene set was curated from Sharp et al. 2019 and analyzed using preranked GSEA as described above.^13^ Heatmaps were generated using pheatmap (v1.0.13), with hierarchical clustering performed using Euclidean distance and complete linkage. All analyses were conducted in R (v4.5.1) using dplyr (v1.1.4), ggplot2 (v3.5.2), msigdbr (v25.1.1), fgsea (v1.34.2), clusterProfiler (v4.16.0), enrichplot (v1.28.4), pheatmap (v1.0.13), ggvenn (v0.1.10), and ggVennDiagram (v1.5.4).

### Immunofluorescence

Cells were fixed with 4% paraformaldehyde (PFA) in phosphate-buffered saline (PBS; pH 7.4) for 10 minutes at room temperature (RT). Following fixation, cells were washed three times with ice-cold PBS (5 minutes per wash). For intracellular targets, cells were permeabilized with 0.3% Triton X-100 in PBS for 15 minutes at RT, followed by three washes with PBS (5 minutes each). Cells were blocked with 3% bovine serum albumin (BSA) in PBS for 30 minutes at RT to reduce nonspecific binding. Primary antibodies were diluted in 3% BSA in PBS: AR (SC-7305; 1:200; Santa Cruz Biotechnology), AR-V7 (RM7; 1:200; RevMAb Biosciences), p53 (SC-126; 1:200; Santa Cruz Biotechnology), and incubated with samples overnight at 4°C in a humidified chamber. After incubation, cells were washed three times with PBS. Secondary antibodies were diluted in 3% BSA in PBS: Anti-Rabbit Alexa Fluor 647 (Ab150079; 1:1000; Abcam), Anti-Mouse Alexa Fluor 568 (Ab175473; 1:1000; Abcam), and applied to samples for 45 minutes at RT in a humidified chamber, protected from light. Cells were washed three times with PBS (5 minutes each). Nuclei were counterstained using Vectashield Plus Antifade Mounting Media with DAPI (Vector Laboratories). Samples were stored protected from light prior to imaging. Images were visualized on Carl Zeiss LSM 780 laser scanning confocal microscope using a 60x Plan-Apochromat (N.A. 1.4) DIC oil objective or 20x Plan-Apochromat (no immersion medium required). Images were pseudocolored and extracted using Zen software.

### Immunohistochemistry

Tumors were routinely processed and embedded in paraffin and serial 5um sections were stained with H&E and chromogenic immunohistochemistry (IHC) on Leica Biosystems’ BondMax autostainer. For IHC, slides were baked at 60C for one hour prior to staining and stained for Ki67 (Cell Signaling Technology 9027, rabbit monoclonal antibody, 1:200) and CyclinD1 (Cell Signaling Technology 2978, rabbit monoclonal, 1:200) using heat-induced epitope retrieval in citrate buffer. Positive tissue controls included human breast cancer for CyclinD1 and human tonsil for CD3. IHC isotype negative controls involved replacing primary antibody with non-clonal, isotype-matched antibody from the same species as the primary antibody. Slides were digitalized at 20× objective (0.5 × 0.5µm per pixel) using Aperio AT2 scanner (Leica Biosystems) and analyzed using HALO (Indica Labs, v4.2). Appropriateness of staining and regions of interest (ROI) annotation were completed by a board-certified pathologist to include viable tumor and exclude tumor necrosis, non-tumor tissues, and histology artifact, including edge artifact and nonspecific staining. ROI for IHC quantification were selected to include areas showing the highest density and intensity of specific staining while avoiding necrosis, folds, edge artifact, nonspecific staining, and non-tumor tissue. One to three ROIs of standardized size ([0.2–0.5 mm²]) were annotated per sample Quantification of staining was performed using HALO Cytonuclear algorithm, which included identifying stain vectors, optimizing cell detection algorithms, and thresholding chromogenic IHC staining based on positive and negative controls. The percentage of positive cells is reported for each marker.

### RNA-seq cohort analysis

Z-normalized mRNA-sequencing data were obtained from cBioPortal (v6.4.4) for four independent cohorts of metastatic prostate adenocarcinoma patients: DFKZ (n = 324), SU2C/PCF (n = 444), TCGA (n = 501), and UW/FH (n = 176).^53–56^ Expression matrices were reshaped to long format and aggregated at the patient level. Within each cohort, samples were ranked by XPO1 expression and stratified into percentile-based quartiles (Q1-Q4). An AR-V7 activity score was computed for each sample as the summed expression (Z-scores) of genes comprising the AR-V7 target gene signature described by Sharp et al. Differences in AR-V7 scores across XPO1 quartiles were visualized using violin plots with overlaid boxplots. Statistical significance was assessed using the Kruskal-Wallis test followed by Dunn’s post hoc testing with Benjamini-Hochberg correction for multiple comparisons. Overall survival analyses were conducted in the SU2C/PCF cohort. Patients with complete overall survival time and status data were included, and sample-to-patient identifiers were harmonized using cohort metadata. Survival probabilities were estimated using Kaplan-Meier analysis comparing the lowest (Q1) and highest (Q4) XPO1 quartiles, with differences assessed by the log-rank test. Cox proportional hazards models were fitted using scaled continuous XPO1 expression (Z-scores) to estimate hazard ratios and corresponding p-values. Tick marks indicate censored observations, and numbers at risk were calculated at defined time intervals. All analyses were performed in R using the dplyr, survival, ggplot2, and ggfortify packages.

## QUANTIFICATION AND STATISTICAL ANALYSIS

All data quantification and figure generation were performed using the software described in the Methods, and final figures were assembled in Adobe Illustrator. Detailed statistical information for each experiment, including the type of test, exact sample size (n), and measures of central tendency and variability (mean ± SD or SEM, or confidence intervals), is provided in the figure legends. Statistical significance is indicated as follows: *p < 0.05, **p < 0.01, ***p < 0.001, and ****p < 0.0001 for all figures.

## DATA AVAILABILITY

All data supporting the conclusions of this study are within the manuscript and/or the Supplementary Information. Global proteomics data have been deposited in the MassIVE repository under accession number MSV000100997 and are publicly available as of the date of publication. This paper does not report original code. Any additional information required to reanalyze the data reported in this paper is available from the lead contact upon request. Requests for further information and resources should be directed to and will be fulfilled by the lead contact, William Douglas Figg, Sr.

## ACKNOWLEDGMENTS

This research was supported by the National Institutes of Health Intramural Research Programs of the National Cancer Institute, Center for Cancer Research project number (ZIA BC 010547) and the National Center for Advancing Translational Sciences (ZIA TR 000047). This project has also been funding in whole or in part with Federal funds from the National Cancer Institute, National Institute of Health, under Contract No. HHSN26120150003I. The contributions of the NIH author(s) were made as part of their official duties as NIH federal employees, are in compliance with agency policy requirements, and are considered Works of the United States Government. However, the findings and conclusions presented in this paper are those of the author(s) and do not necessarily reflect the views of the NIH or the U.S. Department of Health and Human Services, nor does mention of trade names, commercial products, or organizations imply endorsement by the U.S. government.

We thank Drs. Thorkell Andresson and Sudipto Das of the Protein and Metabolite Characterization Core, Cancer Research Technology Program, Frederick National Laboratory for Cancer Research (FNLCR), NIH, for their assistance with proteomic sample processing and mass spectrometry. We also thank Jennifer Matta and Brad Gouker of the Molecular Histopathology Laboratory, Laboratory Animal Sciences Program, FNLCR, NIH, for their assistance with tissue fixation, embedding, sectioning, immunohistochemistry, and histopathologic analysis. We thank Dr. Susan Bates of Columbia University and Dr. Daniel Romo of Baylor University for kindly providing C5-desmethyl PatA compound. We also thank Dr. John Porco of Boston University for kindly providing CR-1-31-B compound.

## AUTHOR CONTRIBUTIONS

Conceptualization, J.D.K, C.H.C., C.J.T., and W.D.F.; methodology, J.D.K, K.B., X.Z., E.L.B., R.D., B.C.B., K.M.W., S.W., C.M., E.B., C.K., R.L., E.E., M.C., C.H.C., C.J.T., and W.D.F.; investigation, J.D.K, K.B., X.Z., E.L.B., S.S.G., R.D., P.J.S., B.C.B., J.L.H., P.S.W., and J.M.C.; writing—original draft, J.D.K, K.B., S.S.G., and C.H.C.; writing—review & editing, all authors; funding acquisition, C.J.T., and W.D.F.; resources, C.J.T., and W.D.F.; supervision, C.H.C., C.J.T., and W.D.F.

## SUPPLEMENTAL INFORMATION

**Document S1. Figures S1-S7**

**Table S1. Single agent drug screening, related to Figure 1**

**Table S2. 42-drug combination screening, related to Figure 2**

**Table S3. Targeted combination drug screening, related to Figure 2**

**Table S4. Apoptosis combination drug screening, related to Figure 3**

**Table S5. Proteomics data, related to Figure 4**

**Table S6. Proteomics significantly up- and downregulated proteins, related to Figure 4**

**Table S7. LNCaP-95 CDX toxicity data, related to Figure 6**

## REFERENCES

1 Freedland, S. J., Davis, M., Epstein, A. J., Arondekar, B. & Ivanova, J. I. Real-world treatment patterns and overall survival among men with Metastatic Castration-Resistant Prostate Cancer (mCRPC) in the US Medicare population. Prostate Cancer Prostatic Dis 27, 327–333 (2024). 10.1038/s41391-023-00725-8

2 Spratt, D. E. et al. Prostate Cancer, Version 3.2026, NCCN Clinical Practice Guidelines In Oncology. J Natl Compr Canc Netw 23, 469–493 (2025). 10.6004/jnccn.2025.0052

3 Antonarakis, E. S. et al. AR-V7 and resistance to enzalutamide and abiraterone in prostate cancer. N Engl J Med 371, 1028–1038 (2014). 10.1056/NEJMoa1315815

4 Beltran, H. et al. The Role of Lineage Plasticity in Prostate Cancer Therapy Resistance. Clin Cancer Res 25, 6916–6924 (2019). 10.1158/1078-0432.CCR-19-1423

5 Beltran, H. et al. Divergent clonal evolution of castration-resistant neuroendocrine prostate cancer. Nat Med 22, 298–305 (2016). 10.1038/nm.4045

6 Ozay, Z. I., Swami, U. & Agarwal, N. Challenges in the Management of Advanced Prostate Cancer. JCO Oncol Pract 21, 447–449 (2025). 10.1200/OP-25-00037

7 Nguyen, M. D., Vinh-Hung, V., Verschraegen, C. & Wang, P. Rethinking combination strategies in metastatic castration-resistant prostate cancer (mCRPC)-lessons from KEYNOTE-641. Transl Androl Urol 15, 5 (2026). 10.21037/tau-2025-aw-747

8 Vincent, F. et al. Phenotypic drug discovery: recent successes, lessons learned and new directions. Nat Rev Drug Discov 21, 899–914 (2022). 10.1038/s41573-022-00472-w

9 Ceribelli, M. et al. Multi-Component, Time-Course screening to develop combination cancer therapies based on synergistic toxicity. Proc Natl Acad Sci U S A 121, e2413372121 (2024). 10.1073/pnas.2413372121

10 Mathews Griner, L. A., et al. High-throughput combinatorial screening identifies drugs that cooperate with ibrutinib to kill activated B-cell-like diffuse large B-cell lymphoma cells. Proc Natl Acad Sci U S A 111, 2349–2354 (2014). 10.1073/pnas.1311846111

11 Lin, G. L. et al. Therapeutic strategies for diffuse midline glioma from high-throughput combination drug screening. Sci Transl Med 11 (2019). 10.1126/scitranslmed.aaw0064

12 Aboukameel, A. et al. Down-regulation of AR splice variants through XPO1 suppression contributes to the inhibition of prostate cancer progression. Oncotarget 9, 35327–35342 (2018). 10.18632/oncotarget.26239

13 Sharp, A. et al. Androgen receptor splice variant-7 expression emerges with castration resistance in prostate cancer. J Clin Invest 129, 192–208 (2019). 10.1172/JCI122819

14 Otte, K. et al. Eltanexor Effectively Reduces Viability of Glioblastoma and Glioblastoma Stem-Like Cells at Nano-Molar Concentrations and Sensitizes to Radiotherapy and Temozolomide. Biomedicines 10 (2022). 10.3390/biomedicines10092145

15 Rosen, E. et al. Phase 1/2 dose expansion study evaluating first-in-class eIF4A inhibitor zotatifin in patients with ER+ metastatic breast cancer. Journal of Clinical Oncology 41, 1080–1080 (2023). 10.1200/JCO.2023.41.16_suppl.1080

16 Meric-Bernstam, F. et al. First-in-human phase 1/2 dose escalation and expansion study evaluating first-in-class eIF4A inhibitor zotatifin in patients with solid tumors. Journal of Clinical Oncology 40, 3081–3081 (2022). 10.1200/JCO.2022.40.16_suppl.3081

17 Nair, A. B. & Jacob, S. A simple practice guide for dose conversion between animals and human. J Basic Clin Pharm 7, 27–31 (2016). 10.4103/0976-0105.177703

18 Cornell, R. F. et al. A phase 1 clinical trial of oral eltanexor in patients with relapsed or refractory multiple myeloma. Am J Hematol 97, E54–E58 (2022). 10.1002/ajh.26420

19 Lee, S. et al. Oral eltanexor treatment of patients with higher-risk myelodysplastic syndrome refractory to hypomethylating agents. J Hematol Oncol 15, 103 (2022). 10.1186/s13045-022-01319-y

20 Gerson-Gurwitz, A. et al. Zotatifin, an eIF4A-Selective Inhibitor, Blocks Tumor Growth in Receptor Tyrosine Kinase Driven Tumors. Front Oncol 11, 766298 (2021). 10.3389/fonc.2021.766298

21 Pianigiani, G. et al. Prolonged XPO1 inhibition is essential for optimal antileukemic activity in NPM1-mutated AML. Blood Adv 6, 5938–5949 (2022). 10.1182/bloodadvances.2022007563

22 Etchin, J. et al. KPT-8602, a second-generation inhibitor of XPO1-mediated nuclear export, is well tolerated and highly active against AML blasts and leukemia-initiating cells. Leukemia 31, 143–150 (2017). 10.1038/leu.2016.145

23 Hing, Z. A. et al. Next-generation XPO1 inhibitor shows improved efficacy and in vivo tolerability in hematological malignancies. Leukemia 30, 2364–2372 (2016). 10.1038/leu.2016.136

24 Ishizawa, J., Kojima, K., Hail, N., Jr., Tabe, Y. & Andreeff, M. Expression, function, and targeting of the nuclear exporter chromosome region maintenance 1 (CRM1) protein. Pharmacol Ther 153, 25–35 (2015). 10.1016/j.pharmthera.2015.06.001

25 Quintanal-Villalonga, A. et al. Exportin 1 inhibition prevents neuroendocrine transformation through SOX2 down-regulation in lung and prostate cancers. Sci Transl Med 15, eadf7006 (2023). 10.1126/scitranslmed.adf7006

26 Gravina, G. L. et al. KPT-330, a potent and selective exportin-1 (XPO-1) inhibitor, shows antitumor effects modulating the expression of cyclin D1 and survivin [corrected] in prostate cancer models. BMC Cancer 15, 941 (2015). 10.1186/s12885-015-1936-z

27 Wei, X. X. et al. A Phase II Trial of Selinexor, an Oral Selective Inhibitor of Nuclear Export Compound, in Abiraterone- and/or Enzalutamide-Refractory Metastatic Castration-Resistant Prostate Cancer. Oncologist 23, 656–e664 (2018). 10.1634/theoncologist.2017-0624

28 Nachmias, B. & Schimmer, A. D. Targeting nuclear import and export in hematological malignancies. Leukemia 34, 2875–2886 (2020). 10.1038/s41375-020-0958-y

29 Kim, W. K. et al. Androgen deprivation induces double-null prostate cancer via aberrant nuclear export and ribosomal biogenesis through HGF and Wnt activation. Nat Commun 15, 1231 (2024). 10.1038/s41467-024-45489-4

30 Uddin, M. H. et al. Nuclear Export Inhibitor KPT-8602 Synergizes with PARP Inhibitors in Escalating Apoptosis in Castration Resistant Cancer Cells. Int J Mol Sci 22 (2021). 10.3390/ijms22136676

31 Wang, C. et al. Epigenetic regulation of EIF4A1 through DNA methylation and an oncogenic role of eIF4A1 through BRD2 signaling in prostate cancer. Oncogene 41, 2778–2785 (2022). 10.1038/s41388-022-02272-3

32 Wolfe, A. L. et al. RNA G-quadruplexes cause eIF4A-dependent oncogene translation in cancer. Nature 513, 65–70 (2014). 10.1038/nature13485

33 Rubio, C. A. et al. Transcriptome-wide characterization of the eIF4A signature highlights plasticity in translation regulation. Genome Biol 15, 476 (2014). 10.1186/s13059-014-0476-1

34 Kuzuoglu-Ozturk, D. et al. Small-molecule RNA therapeutics to target prostate cancer. Cancer Cell 43, 841–855 e848 (2025). 10.1016/j.ccell.2025.02.027

35 Coleman, R. L., Kalyanapu, P., Walker, C. J. & Vergote, I. An evaluation of selinexor as a maintenance therapy for patients with p53 wild-type, advanced, or recurrent endometrial carcinoma. Expert Rev Anticancer Ther 25, 1007–1019 (2025). 10.1080/14737140.2025.2522948

36 Li, Y. L., Tian, H., Jiang, J., Zhang, Y. & Qi, X. W. Multifaceted regulation and functions of fatty acid desaturase 2 in human cancers. Am J Cancer Res 10, 4098–4111 (2020).

37 Wambach, M. et al. Clinical implications of AGR2 in primary prostate cancer: Results from a large-scale study. APMIS 132, 256–266 (2024). 10.1111/apm.13382

38 Li, J. et al. Arginase 2 is a Diagnostic and Prognostic Marker for Prostate Cancer and Is Associated with Metabolism. Altern Ther Health Med 30, 102–111 (2024).

39 Li, S. et al. TFAP4/DLGAP5 promotes tumor progression and macrophage M2 polarization in prostate cancer by activating the JAK2/STAT3 signaling. Exp Cell Res 452, 114753 (2025). 10.1016/j.yexcr.2025.114753

40 Gu, Y., Jiang, J. & Liang, C. TFAP4 promotes the growth of prostate cancer cells by upregulating FOXK1. Exp Ther Med 22, 1299 (2021). 10.3892/etm.2021.10734

41 Cordova, R. A. et al. GCN2 eIF2 kinase promotes prostate cancer by maintaining amino acid homeostasis. Elife 11 (2022). 10.7554/eLife.81083

42 Subramanian, A. et al. Gene set enrichment analysis: a knowledge-based approach for interpreting genome-wide expression profiles. Proc Natl Acad Sci U S A 102, 15545–15550 (2005). 10.1073/pnas.0506580102

43 Ritz, C., Baty, F., Streibig, J. C. & Gerhard, D. Dose-Response Analysis Using R. PLoS One 10, e0146021 (2015). 10.1371/journal.pone.0146021

44 Gu, Z., Gu, L., Eils, R., Schlesner, M. & Brors, B. circlize Implements and enhances circular visualization in R. Bioinformatics 30, 2811–2812 (2014). 10.1093/bioinformatics/btu393

45 Zheng, S. et al. SynergyFinder Plus: Toward Better Interpretation and Annotation of Drug Combination Screening Datasets. Genomics Proteomics Bioinformatics 20, 587–596 (2022). 10.1016/j.gpb.2022.01.004

46 Beshiri, M. L. et al. A PDX/Organoid Biobank of Advanced Prostate Cancers Captures Genomic and Phenotypic Heterogeneity for Disease Modeling and Therapeutic Screening. Clin Cancer Res 24, 4332–4345 (2018). 10.1158/1078-0432.CCR-18-0409

47 Nguyen, H. M. et al. LuCaP Prostate Cancer Patient-Derived Xenografts Reflect the Molecular Heterogeneity of Advanced Disease an--d Serve as Models for Evaluating Cancer Therapeutics. Prostate 77, 654–671 (2017). 10.1002/pros.23313

48 Li, C. et al. Molecular decoupling of lineage identity and morphology in aggressive variant prostate cancer. medRxiv (2026). 10.64898/2026.01.07.26343520

49 Brim, B. C. et al. Direct Co-Targeting of Bcl-xL and Mcl-1 Exhibits Synergistic Effects in AR-V7-Expressing CRPC Models. Cancer Res Commun 5, 1396–1408 (2025). 10.1158/2767-9764.CRC-25-0096

50 Korotkevich, G. S., V.; Budin, N.; Shpak, B.; Artyomov, M. N.; Sergushichev, A. Fast preranked gene set enrichment analysis using adaptive multilevel splitting. bioRxiv (2021). 10.1101/060012

51 Liberzon, A. et al. The Molecular Signatures Database (MSigDB) hallmark gene set collection. Cell Syst 1, 417–425 (2015). 10.1016/j.cels.2015.12.004

52 Yu, G., Wang, L. G., Han, Y. & He, Q. Y. clusterProfiler: an R package for comparing biological themes among gene clusters. OMICS 16, 284–287 (2012). 10.1089/omi.2011.0118

53. Cancer Genome Atlas Research, N. The Molecular Taxonomy of Primary Prostate Cancer. Cell 163, 1011–1025 (2015). 10.1016/j.cell.2015.10.025

54 Abida, W. et al. Genomic correlates of clinical outcome in advanced prostate cancer. Proc Natl Acad Sci U S A 116, 11428–11436 (2019). 10.1073/pnas.1902651116

55 Kumar, A. et al. Substantial interindividual and limited intraindividual genomic diversity among tumors from men with metastatic prostate cancer. Nat Med 22, 369–378 (2016). 10.1038/nm.4053

56 Gerhauser, C. et al. Molecular Evolution of Early-Onset Prostate Cancer Identifies Molecular Risk Markers and Clinical Trajectories. Cancer Cell 34, 996–1011 e1018 (2018). 10.1016/j.ccell.2018.10.016

