## Supplementary Figure S1-S7 for "Co-Targeting Nuclear Export and Translation Initiation Uncovers a Therapeutic Vulnerability in Lethal Prostate Cancer"

Running Title: XPO1-EIF4A1 co-targeting in prostate cancer

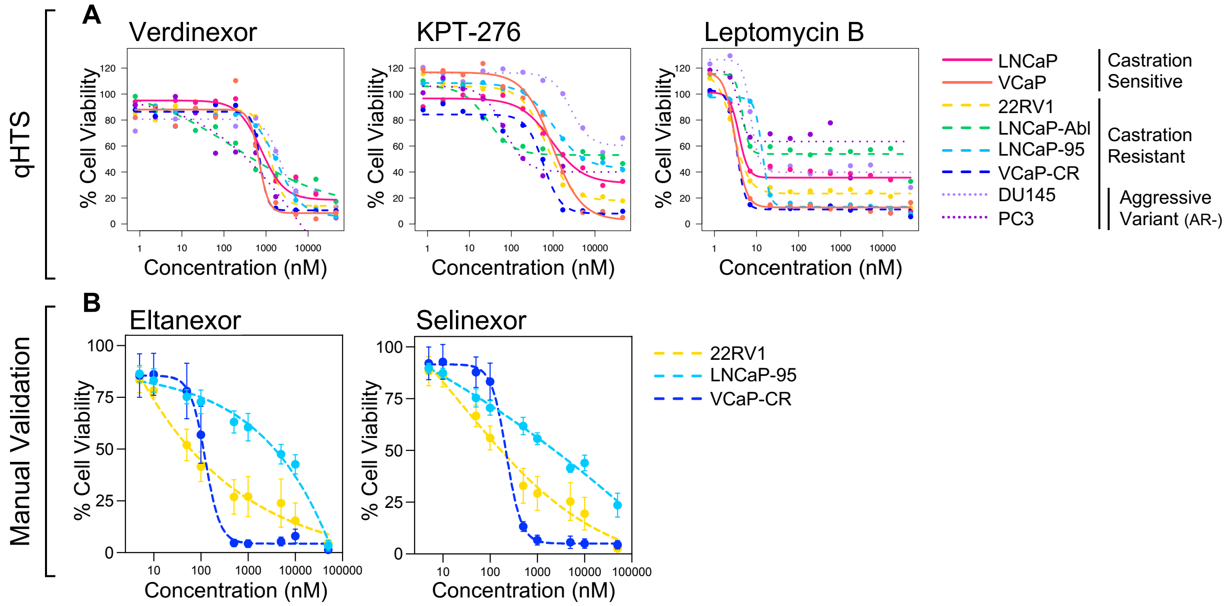


**Figure S1. Single agent activity of XPO1 inhibitors in prostate cancer models, related to Figure 1.**

(A) Single agent dose-response curves displaying percent cell viability following treatment with the XPO1 inhibitors Verdinexor, KPT-276, and Leptomycin B across an 11-point concentration range (0.8 nM-45 μM; 1:3 dilution series), assessed by CellTiter-Glo (CTG).

(B) Manual validation of cell viability following treatment with Eltanexor or Selinexor in AR-V7-positive 22RV1, LNCaP-95, and VCaP-CR cells, assessed by CTG at 48 h (22RV1, LNCaP-95) or 72 h (VCaP-CR). Data represent mean ± SD from n=3 replicates.

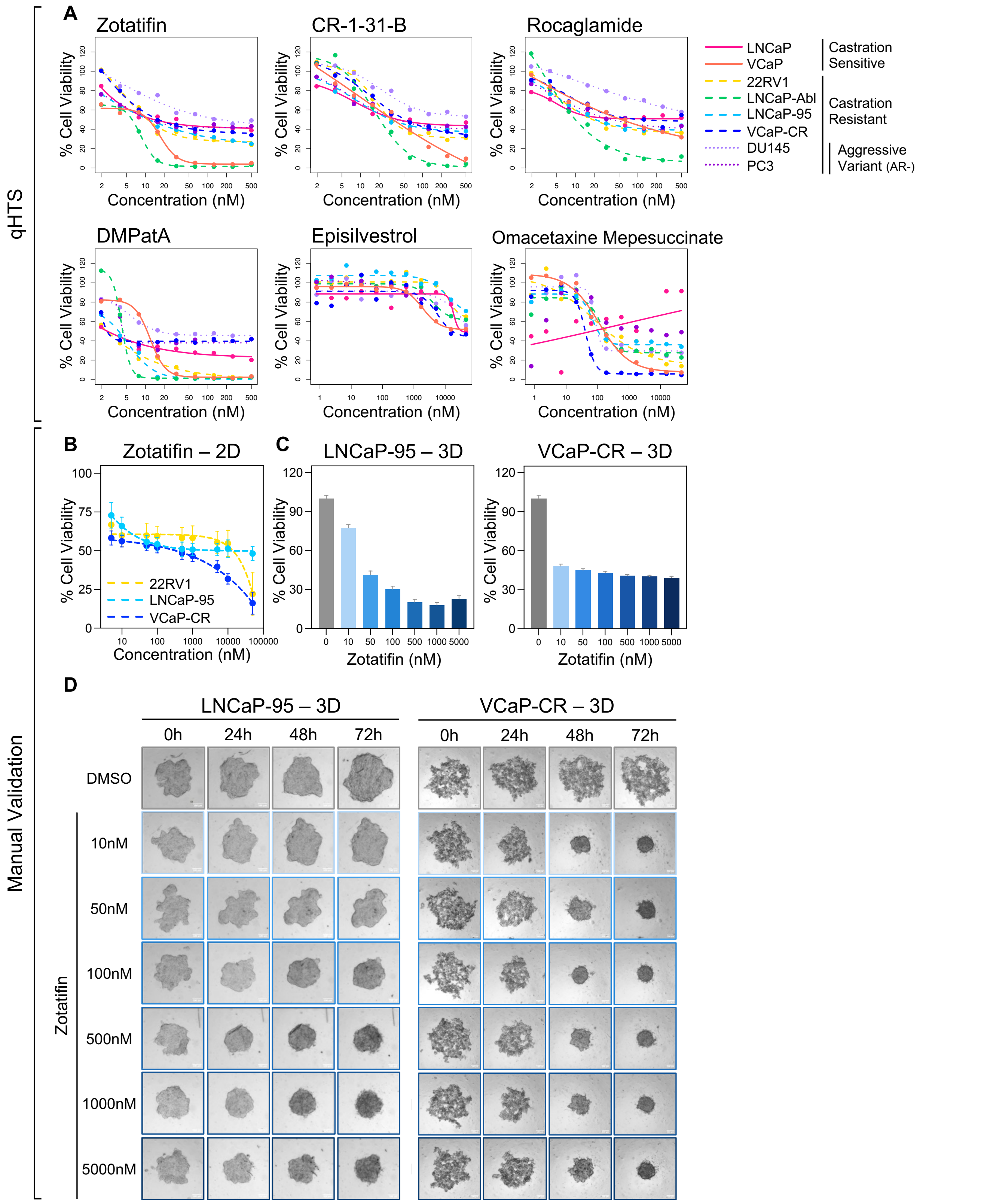


**Figure S2. Single agent activity of EIF4A1 inhibitors in prostate cancer models, related to Figure 2.**

(A) Single agent dose-response curves displaying percent cell viability following treatment with the EIF4A1 inhibitors Zotatifin, CR-1-31-B, Rocaglamide, DMPatA, Episilvestrol, and Omacetaxine mepesuccinate across an 11-point concentration range (0.8 nM-45 μM; 1:3 dilution series), assessed by CTG.

(B) Manual validation of cell viability following treatment with Zotatifin in AR-V7-positive 22RV1, LNCaP-95, and VCaP-CR cells, assessed by CTG at 48 h (22RV1, LNCaP-95) or 72 h (VCaP-CR). Data represent mean ± SD from n=3 replicates.

(C) Manual validation of cell viability in 3D LNCaP-95 spheroids and 3D VCaP-CR cell aggregates following treatment with Zotatifin, assessed by 3D CTG at 48 h (LNCaP-95) or 72 h (VCaP-CR). Data represent mean ± SEM from n=3 replicates.

(D) Representative brightfield images of LNCaP-95 3D spheroids and VCaP-CR 3D cell aggregates treated with Zotatifin for 0, 24, 48, and 72 hours. Scale bar, 100 µm.

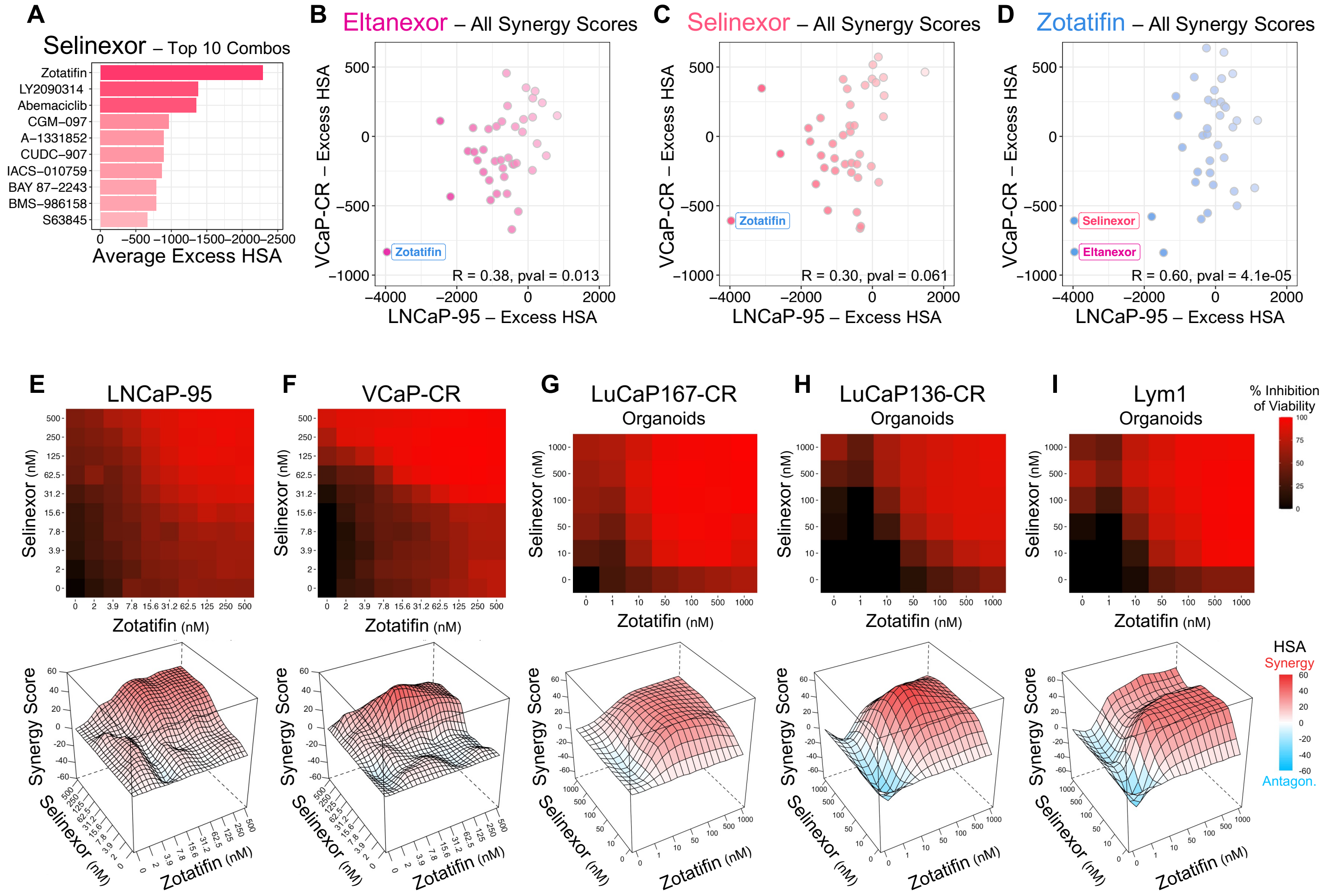


**Figure S3. Combination screening reveals synergy between XPO1 and EIF4A1 inhibition, related to Figure 2.**

(A) Top 10 most synergistic combinations with Selinexor, ranked by mean Excess HSA across LNCaP-95 and VCaP-CR cells.

(B) Scatter plot comparing Excess HSA synergy scores for combinations with Eltanexor across VCaP-CR and LNCaP-95 cells from the all-versus-all combination screen. Each point represents a distinct drug combination; Pearson correlation coefficient (R) and p-value are shown.

(C) As in (B), but for combinations with Selinexor as the anchor compound.

(D) As in (B), but for combinations with Zotatifin as the anchor compound; XPO1 inhibitors (Eltanexor and Selinexor) are highlighted.

(E-F) Focused 10 × 10 dose-response matrices for the Selinexor + Zotatifin combination extracted from the screen and analyzed using SynergyFinder in (E) LNCaP-95 and (F) VCaP-CR cells. Heatmaps show percent inhibition of cell viability, with corresponding three-dimensional surface plots depicting HSA synergy scores.

(G-I) Validation of Selinexor + Zotatifin synergy in patient-derived organoid models. Percent inhibition heatmaps and three-dimensional HSA synergy surface plots are shown for (G) LuCaP167-CR, (H) LuCaP136-CR, and (I) Lym1 organoids. Data represent the mean from n=3 replicates.

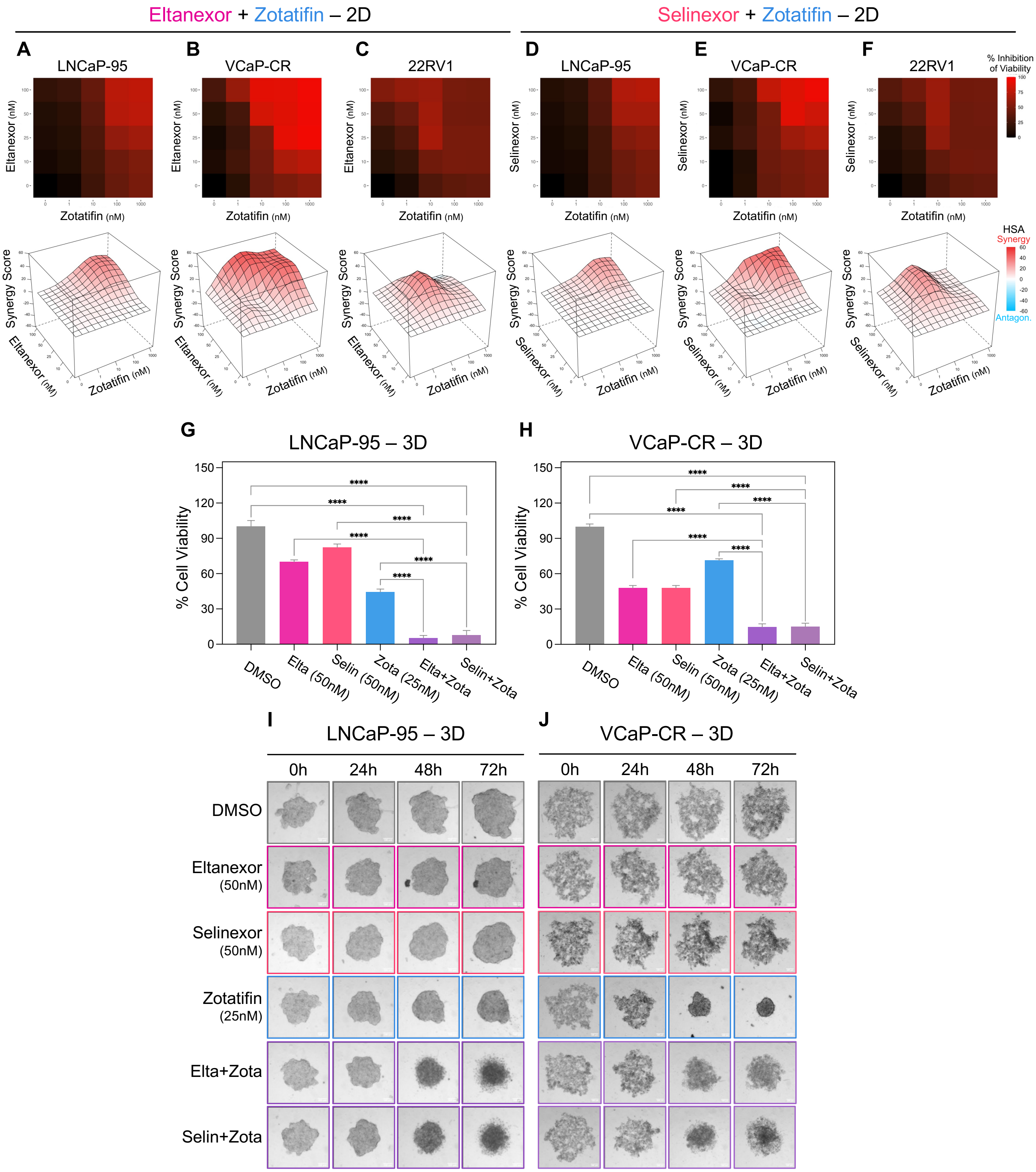


**Figure S4. Manual validation of synergy between XPO1 and EIF4A1 inhibition, related to Figure 2.** (A-C) Cell viability following combination treatment was measured in 96-well plates for (A) LNCaP-95, (B) VCaP-CR, and (C) 22RV1 cells grown in 2D culture. Viability was assessed by CTG at 48 h (LNCaP-95, 22RV1) or 72 h (VCaP-CR). Heatmaps display percent inhibition, with corresponding three-dimensional

Excess HSA synergy surfaces shown. Data represent the mean from n=3 replicates.

(D-F) As in (A-C), for Selinexor + Zotatifin combinations.

(G-H) LNCaP-95 3D spheroids (G, 48 h) and VCaP-CR 3D aggregates (H, 72 h) treated with DMSO, single agents, or combinations; bar plots show percent viability relative to control. Data represent mean ± SEM from n=3 replicates. Combination effects were compared to vehicle or single agents by one-way ANOVA with multiple-comparisons testing; ****$p < 0.0001$.

(I-J) Representative brightfield images of LNCaP-95 spheroids (I) and VCaP-CR aggregates (J) at 0, 24, 48, and 72 h following indicated treatment. Scale bar, 100 μm.

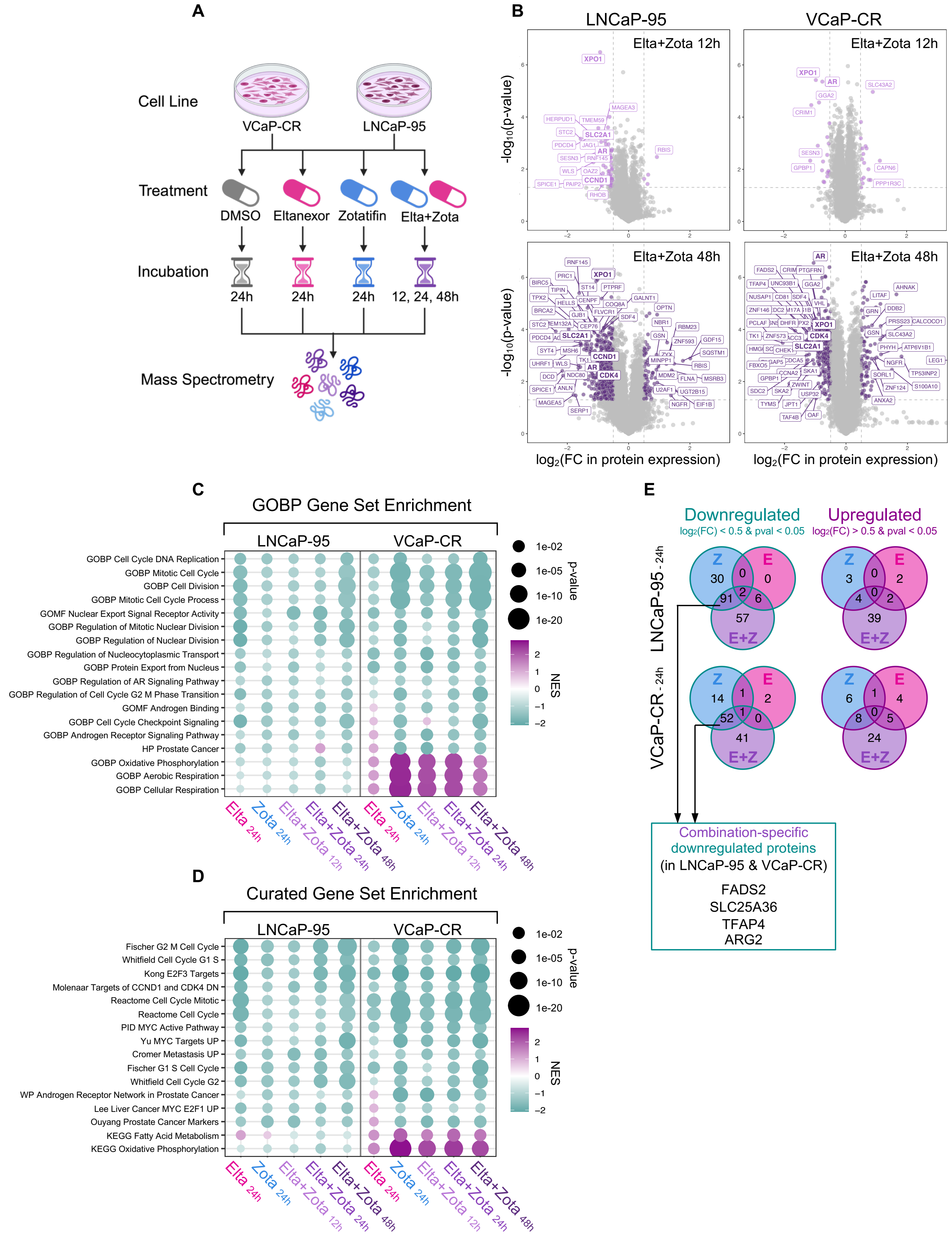

A
Cell Line
VCaP-CR
LNCaP-95
Treatment
DMSO
Eltanexor
Zotatifin
Elta+Zota
Incubation
24h
24h
24h
12, 24, 48h
Mass Spectrometry
B
LNCaP-95
VCaP-CR
Elta+Zota 12h
Elta+Zota 48h
-log10(p-value)
log2(FC in protein expression)
XPO1
AR
SLC2A1
CCND1
CDK4
C
GOBP Gene Set Enrichment
LNCaP-95
VCaP-CR
GOBP Cell Cycle DNA Replication
GOBP Mitotic Cell Cycle
GOBP Cell Division
GOBP Mitotic Cell Cycle Process
GOMF Nuclear Export Signal Receptor Activity
GOBP Regulation of Mitotic Nuclear Division
GOBP Regulation of Nuclear Division
GOBP Regulation of Nucleocytoplasmic Transport
GOBP Protein Export from Nucleus
GOBP Regulation of AR Signaling Pathway
GOBP Regulation of Cell Cycle G2 M Phase Transition
GOMF Androgen Binding
GOBP Cell Cycle Checkpoint Signaling
GOBP Androgen Receptor Signaling Pathway
HP Prostate Cancer
GOBP Oxidative Phosphorylation
GOBP Aerobic Respiration
GOBP Cellular Respiration
p-value
1e-02
1e-05
1e-10
1e-20
NES
Elta 24h
Zota 24h
Elta+Zota 12h
Elta+Zota 24h
Elta+Zota 48h
D
Curated Gene Set Enrichment
Fischer G2 M Cell Cycle
Whitfield Cell Cycle G1 S
Kong E2F3 Targets
Molenaar Targets of CCND1 and CDK4 DN
Reactome Cell Cycle Mitotic
Reactome Cell Cycle
PID MYC Active Pathway
Yu MYC Targets UP
Cromer Metastasis UP
Fischer G1 S Cell Cycle
Whitfield Cell Cycle G2
WP Androgen Receptor Network in Prostate Cancer
Lee Liver Cancer MYC E2F1 UP
Ouyang Prostate Cancer Markers
KEGG Fatty Acid Metabolism
KEGG Oxidative Phosphorylation
E
Downregulated
log2(FC) < 0.5 & pval < 0.05
Upregulated
log2(FC) > 0.5 & pval < 0.05
LNCaP-95 - 24h
VCaP-CR - 24h
Z
E
E+Z
Combination-specific downregulated proteins (in LNCaP-95 & VCaP-CR)
FADS2
SLC25A36
TFAP4
ARG2

**Figure S5. Extended proteomic analysis of XPO1 and EIF4A1 co-inhibition, related to Figure 4.**
(A) Experimental schematic showing LNCaP-95 and VCaP-CR cells treated with DMSO or single agents for 24 h. Combination-treated samples were analyzed at 12, 24, and 48 h. Created with Biorender.com.
(B) Global quantitative proteomic analysis of LNCaP-95 and VCaP-CR cells treated with Eltanexor (50 nM), Zotatifin (LNCaP-95: 25 nM; VCaP-CR: 10 nM), or the Eltanexor + Zotatifin combination for 12 or 48 hours. Data represent the mean from n=3 replicates. Protein abundance changes were assessed relative to DMSO control, and results are displayed as volcano plots depicting $\log_2$(fold-change) versus $-\log_{10}$(p-value). Selected significantly upregulated and downregulated proteins are highlighted.
(C) MSigDB C5: Gene Ontology Biological Process (GOBP) pathway enrichment analysis of LNCaP-95 and VCaP-CR proteomic datasets, ranked by differential protein abundance for each treatment condition relative to DMSO. Dot plots display normalized enrichment scores (NES) for significantly enriched pathways, with dot size indicating statistical significance.
(D) As in (C), selected enrichment analysis of MSigDB C2: curated gene sets (including BioCarta, KEGG, PID, Reactome, and WikiPathways).
(E) Venn diagrams depicting proteins significantly downregulated (left) or upregulated (right) by 24 h treatment of Zotatifin (Z), Eltanexor (E), or the combination (E+Z) in LNCaP-95 (top) and VCaP-CR (bottom) cells.

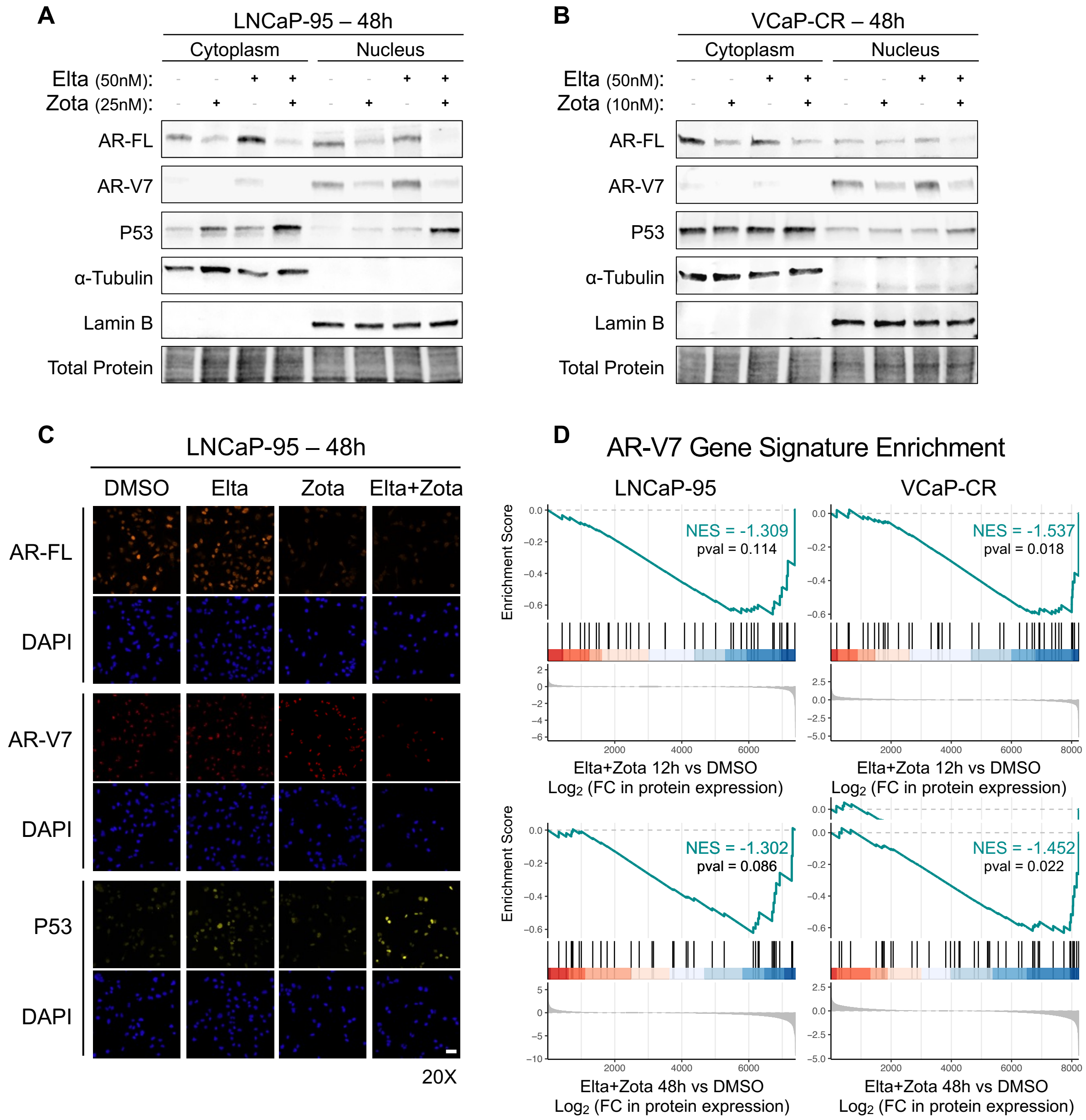


**Figure S6. Extended analysis of AR/AR-V7 signaling and p53 regulation following combined XPO1 and EIF4A1 inhibition, related to Figure 5.**

(A-B) Subcellular fractionation followed by immunoblot analysis of cytoplasmic and nuclear extracts from (A) LNCaP-95 and (B) VCaP-CR cells treated with Eltanexor, Zotatifin, or the combination for 48 hours. Total protein staining is shown as a loading control.

(C) Immunofluorescence staining of LNCaP-95 cells treated with Eltanexor, Zotatifin, or the combination for 48 hours, acquired at 20× magnification; nuclei are marked by DAPI. Scale bar, 50 µm.

(D) Gene set enrichment analysis of the AR-V7 gene signature performed on global proteomics datasets following combination treatment for 12 or 48 hours relative to DMSO in LNCaP-95 and VCaP-CR cells.

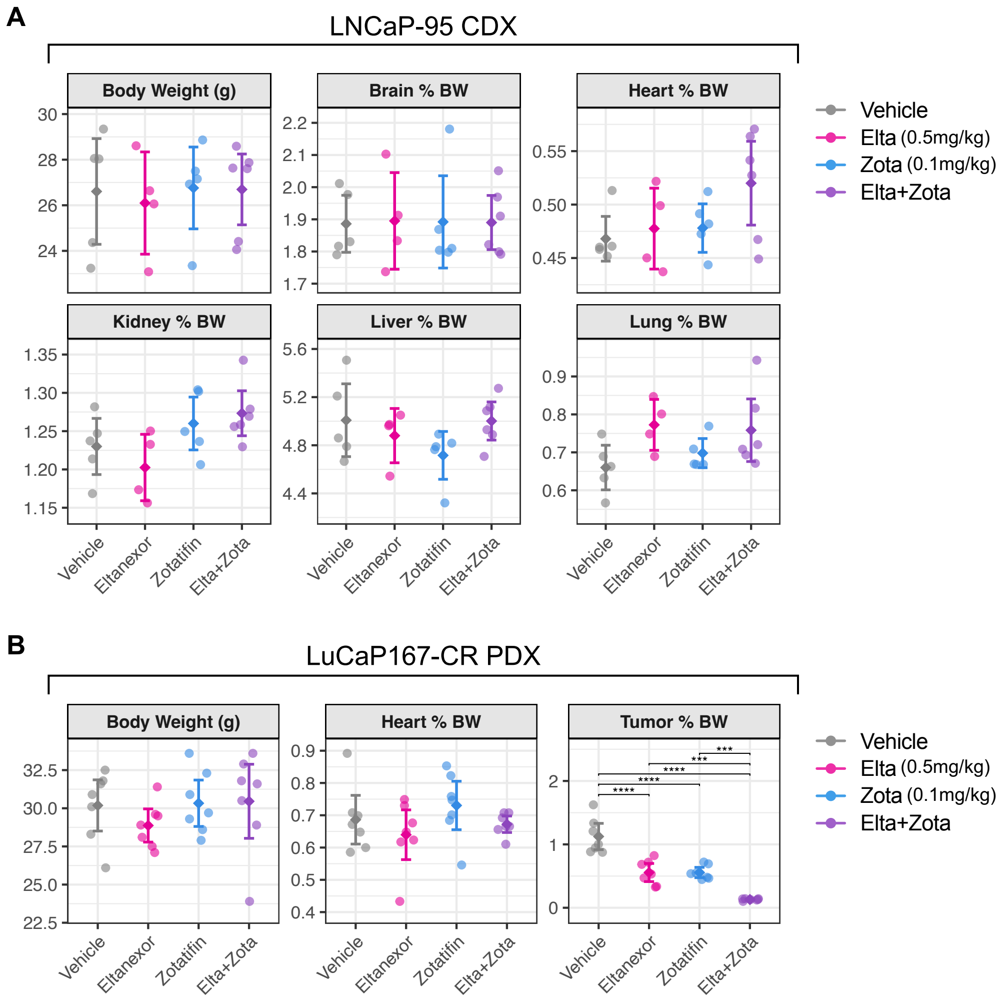


**Figure S7. Systemic tolerability of combined XPO1 and EIF4A1 inhibition in vivo, related to Figure 6.**
(A) Body weight (g) and organ (brain, heart, kidney, liver, and lung) weights normalized to body weight (BW) in LNCaP-95 cell-derived xenograft (CDX) mice (n=5 per treatment group) at the end of 3 weeks of treatment with vehicle, Eltanexor, Zotatifin, or the combination.
(B) Body weight, heart weight (% BW), and tumor weight (% BW) in LuCaP167-CR patient-derived xenograft (PDX) mice (n=7 per treatment group) at the end of 4 weeks of treatment with vehicle, Eltanexor, Zotatifin, or the combination. Points represent individual mice; bars indicate mean ± 95% confidence interval. Statistical significance was assessed by one-way ANOVA followed by Tukey's multiple-comparisons test. Significance is indicated as ***$p < 0.001$, ****$p < 0.0001$.
